# DAP5 enables translation re-initiation on structured messenger RNAs

**DOI:** 10.1101/2021.01.21.427569

**Authors:** Ramona Weber, Leon Kleemann, Insa Hirschberg, Min-Yi Chung, Eugene Valkov, Cátia Igreja

## Abstract

Half of mammalian transcripts contain short upstream open reading frames (uORFs) that potentially regulate translation of the downstream coding sequence (CDS). The molecular mechanisms governing these events remain poorly understood. Here we find that the non-canonical initiation factor Death-associated protein 5 (DAP5 or eIF4G2) is selectively required for re-initiation at the main CDS following uORF translation. Using ribosome profiling and luciferase-based reporters coupled with mutational analysis we show that DAP5-mediated re-initiation occurs on messenger RNAs (mRNAs) with long, structure-prone 5′ leader sequences and persistent uORF translation. These mRNAs preferentially code for signalling factors such as kinases and phosphatases. We also report that cap/eIF4F- and eIF4A-dependent recruitment of DAP5 to the mRNA facilitates re-initiation by unrecycled post-termination 40S subunits. Our study reveals important mechanistic insights into how a non-canonical translation initiation factor involved in stem cell fate shapes the synthesis of specific signalling factors.

## Introduction

Translation initiation directs the ribosome to the start codon of messenger (m)RNAs. Binding of the eukaryotic initiation factor (eIF) 4F complex to the 5′ cap m GpppX structure, present in all eukaryotic mRNAs, triggers the vast majority of translation initiation events (Topisirovic et al., 2011). The eIF4F complex consists of three subunits: eIF4G, eIF4A and eIF4E. As the scaffolding factor of the complex, eIF4G bridges the cap-binding protein eIF4E and the ATP-dependent RNA helicase eIF4A which unwinds secondary structures within the 5′ untranslated region (UTR) of the mRNA (Pelletier and Sonenberg, 2019). In addition, eIF4G associates with the poly(A) binding protein (PABP) to link the 5′ and 3′ ends of the mRNA, and recruits the 43S preinitiation complex (PIC; 40S ribosomal subunit bound to the eIF2:GTP:Met–tRNAi^Met^ ternary complex, eIF3, eIF1 and eIF1A) to capped mRNAs (Hashem and Frank, 2018; Merrick and Pavitt, 2018). PIC scanning along the 5′ UTR to the AUG start codon and joining of the 60S ribosomal subunit determines the assembly of an elongation-committed 80S ribosome (Merrick and Pavitt, 2018).

In mammalian cells, three related eIF4G proteins regulate initiation of translation. Unlike eIF4G1 (hereafter eIF4G) and eIF4G3, Death-associated protein 5 (DAP5, eIF4G2, NAT1 or p97) lacks the eIF4E and PABP-binding sites, but is homologous to the middle domain of eIF4G (MIF4G), the MA3, and the W2 C-terminal domains (Figure 1A). The MIF4G interacts with eIF4A and eIF3 (Imataka et al., 1997), whereas the W2 domain of DAP5 associates with the β subunit of eIF2 (eIF2β) (Lee and McCormick, 2006; Liberman et al., 2015) instead of the MAPK-interacting kinases MNK1 (Pyronnet et al., 1999) and MKN2 (Scheper et al., 2001). Thus, DAP5 is considered to be a non-canonical factor different from the other eIF4G homologs, that directs the ribosome to the mRNA independently of eIF4F.

**Figure 1.**
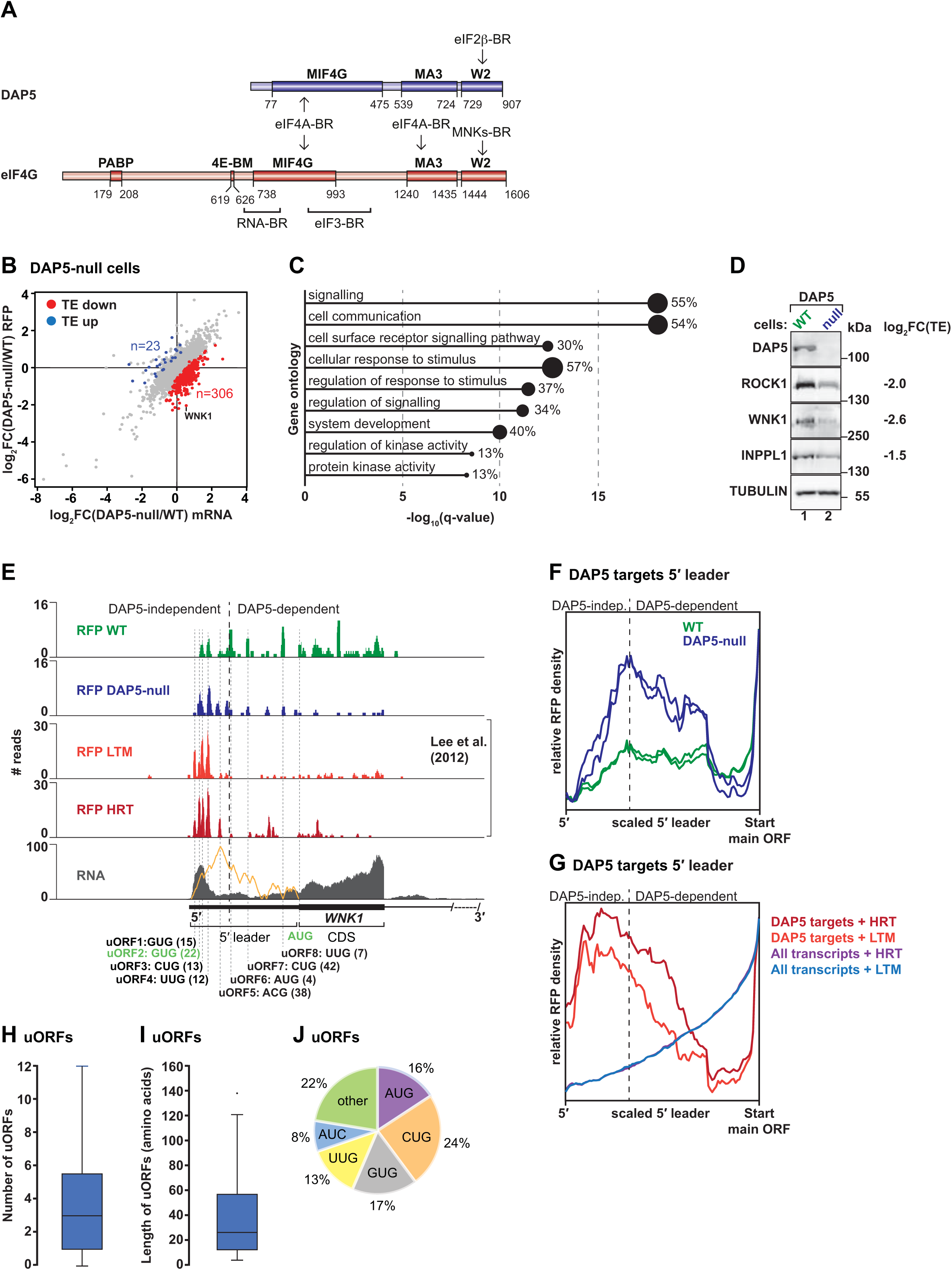
DAP5 mediates the synthesis of signalling proteins. **(A)** Schematic representation of the DAP5 and eIF4G proteins. PABP: poly(A)-binding protein-binding region; 4E-BM: eIF4E-binding motif; MIF4G: middle eIF4G domain; MA3: MA3 domain; W2: W2 domain; eIF4A-BR: eIF4A-binding region; eIF2β-BR: eIF2β-binding region; MNKs-BR: MNK1 and MNK2-binding region; RNA-BR: RNA-binding region; eIF3-BR: eIF3-binding region. The amino acid positions at the domain/motif boundaries are indicated below the proteins. The MIF4G domains of DAP5 and eIF4G, and the MA3 domain of eIF4G bind to eIF4A. The MIF4G domains are also known to contribute to the interactions with RNA and the eIF3 complex. The W2 domain of DAP5 interacts with eIF2β whereas the corresponding domain in eIF4G associates with the MNK kinases 1 and 2. **(B)** Comparative analysis of translation efficiency (TE) in wild type (WT) and DAP5-null HEK293T cells. Genes were plotted as a scatter graph according to changes in ribosome occupancy [log_2_FC RFP] on the y axis and mRNA abundance [log_2_FC mRNA] on the x axis. Each dot represents an individual gene (n_total_=9870) selected using FPKM >2. Genes without changes in TE are indicated in grey. Genes with increased or decreased TE are highlighted in blue (23 genes) and red (306 genes), respectively. See also Figure S1. **(C)** Gene ontology (GO)-terms associated with the genes with decreased TE in cells lacking DAP5. Bar graph shows -log_10_ *q* values for each of the overrepresented category. Values and black circles indicate the % of genes within each category. **(D)** Immunoblotting demonstrating the loss of DAP5 in the null cells. In agreement with the observed changes in TE [log2FC(TE)], ROCK1 and WNK1 kinases, and INPPL1 phosphatase protein levels decrease in the absence of DAP5. TUBULIN served as loading control. Blots were probed with antibodies recognizing DAP5, ROCK1, WNK1, INPPL1 and TUBULIN. **(E)** Ribosome footprints and total mRNA reads distribution along *WNK1* exon 1 including the 5′ leader and the most 5′ proximal coding sequence in WT and DAP5-null cells. Also shown are the ribosome footprint profiles (RFPs) in HEK293 cells treated with harringtonine (HRT) and lactimidomycin (LTM) obtained by Lee and co-workers (Lee et al., 2012). The predicted propensity for secondary structure across *WNK1* 5′ leader, determined using the ViennaRNA package 2.0 (Lorenz et al., 2011), is illustrated in orange in the mRNA panel. uORFs position in the 5′ leader is indicated with the corresponding start codons (GUG, CUG, UUG, ACG or AUG). Start codons highlighted in green are in frame with the AUG at the main annotated coding sequence of *WNK1*. Gene annotation is depicted below the profiles. DAP5-independent and -dependent translation is indicated with a black dashed line. CDS: coding sequence. See also Figure S2. **(F, G)** Metagene analyses of ribosome density at the 5′ leaders of the 306 DAP5 targets in WT (green) and DAP5-null (blue) cells (F), and 5′ leaders of DAP5 targets and all expressed transcripts expressed in HEK293 cells treated with HRT or LTM (Lee et al., 2012) (G). Ribosome densities were determined as the ratio of footprints within the 5′ leader relative to the footprints at the annotated downstream CDS start codon. The black dashed line indicating DAP5-independent (indep.) and DAP5-dependent translation was defined as the position along the 5′ leaders in which RFP density decreases in the absence of DAP5. **(H, I)** Variation of uORF number and length in the 5′ leaders of all DAP5 targets (n=306). Boxes indicate the 25^th^ to 75^th^ percentiles; black line inside the box represents the median; whiskers indicate the extend of the highest and lowest observations; dots show the outliers. **(J)** Start codon usage of uORFs preferentially translated in the absence of DAP5.

Multiple mechanisms have been proposed to explain how DAP5 engages in translation initiation. Most studies indicate that DAP5 stimulates translation using internal ribosome entry sites (IRESes) or cap-independent translation enhancers (CITEs) present in the 5′ UTR of specific mRNAs when cells are exposed to conditions that hinder cap-dependent initiation (Haizel et al., 2020; Henis-Korenblit et al., 2002; Henis-Korenblit et al., 2000; Hundsdoerfer et al., 2005; Lee and McCormick, 2006; Lewis et al., 2008; Liberman et al., 2009; Marash et al., 2008; Weingarten-Gabbay et al., 2014). Alternatively, DAP5 was proposed to initiate non-canonical translation via the assembly of cap-bound complexes with proteins other than eIF4E (Bukhari et al., 2016; de la Parra et al., 2018). As DAP5 is an ubiquitously expressed and abundant protein known to control the expression of genes required for stem cell differentiation and embryonic development (Nousch et al., 2007; Sugiyama et al., 2017; Takahashi et al., 2020; Yamanaka et al., 2000; Yoffe et al., 2016; Yoshikane et al., 2007), understanding its mode of action is important for elucidating the mechanisms that drive non-canonical translation.

Upstream open reading frames (uORFs) are prevalent and translated in the 5′ UTRs (hereafter 5′ leaders) of mammalian mRNAs (Bazzini et al., 2014; Chen et al., 2020; Fritsch et al., 2012; Ingolia, 2014; Ingolia et al., 2011; Lee et al., 2012). Expression of downstream and main coding sequences (CDSes) requires scanning of the PIC past the uORFs (leaky scanning) or re-initiation by unrecycled ribosomal complexes after uORF translation (Jackson et al., 2012). Despite the regulatory roles often attributed to uORFs in gene expression and disease (Barbosa et al., 2013), the mechanisms and proteins involved in uORF and main CDS translation in eukaryotic mRNAs are incompletely understood. Here, we describe DAP5 as a non-canonical factor that enhances translation of capped mRNAs via re-initiation. DAP5 is crucial for the translation of particular CDSes after uORF translation in transcripts with structured 5′ leaders. Together with eIF4A, DAP5 regulates the translation of transcripts encoding signalling and regulatory factors with important roles in stem cell and cancer biology, such as kinases and phosphatases. Our findings reveal an unexpected role for a member of the eIF4G family of proteins in the control of translation in human cells.

## Results

### DAP5 mediates the synthesis of signalling proteins

To study the function of DAP5 in translation, we determined the translational landscape of CRISPR-Cas9 engineered DAP5-null and wild type (WT) HEK293T cells (Figure S1A-C). After isolation and sequencing of ribosome-protected mRNA fragments (ribosome profiling or Ribo-Seq) and matched transcriptome analysis (RNA-Seq), we determined genome-wide transcriptional and translational changes (Figures 1B and S1D) (Ingolia et al., 2011; Zhong et al., 2017). The Ribo-Seq and RNA-Seq experiments were reproducible as replicates clustered together (Figures S1E, F).

In the absence of DAP5, a group of genes – hereafter referred to as DAP5 targets – showed a significant reduction in translation efficiency (TE; ribosome occupancy/mRNA abundance) (Figure 1B; n=306, red). Although the majority of DAP5 target transcripts were more abundant, the number of ribosomes per mRNA (footprints or occupancy) decreased in the null cells (Table S1). Other translatome-associated differences were found in a small cohort of mRNAs with increased TE in the null cells (n=23, blue; Figure 1B and Table S1). In addition to translatome-only changes, we also observed pronounced differences in transcript abundance in the null cells (n=3537; Figure S1D and Table S1). These differences may result from effects on transcription and/or mRNA turnover following DAP5 depletion.

Notably, DAP5 targets included mRNAs encoding proteins involved in cell signalling, or cellular response to stimuli, such as the serine/threonine-protein kinases WNK1 [With-No-Lysine (K)1] and ROCK1 (Rho-associated protein kinase 1), the RAC-alpha serine/threonine-protein kinase AKT1 or the phosphatidylinositol 3,4,5-triphosphate 5-phosphatase 2 (INPPL1, a.k.a. SHIP2), among others (Figure 1C, Table S1). Accordingly, WNK1, ROCK1 and INPPL1 protein levels assessed by immunoblotting were diminished in the absence of DAP5 (Figure 1D). Importantly, decreased protein synthesis in the null cells was not caused by deficiency in the expression of eIF4E, eIF4G, eIF4A and PABP (Figure S1C), or changes in global translation (Figure S1G). With the exception of a reproducible increase in the free 40S subunits peak, polysome profiles of DAP5-null cells after sucrose density gradient separation were similar to those of wild type cells (Figure S1G-I). However, the association of *WNK1* and *ROCK1* mRNAs with polysomes, but not *GAPDH*, shifted from the heavy to the light polysome fractions (Figures S1J-L, lanes 16-18 vs. 12-15) in the absence of DAP5. These results indicate that the translational efficiency of a specific subset of transcripts is regulated by DAP5.

### DAP5 targets 5**′** leaders have unique features

In addition to the differences in TE, we also observed qualitative changes in the pattern of ribosomal occupancies (footprints) in DAP5 target mRNAs. Ribosome occupancy at the annotated (main) CDSes was markedly decreased in the absence of DAP5. Moreover, ribosome footprints were skewed towards the 5′ leaders of these transcripts (Figures 1E and S2). Estimation of footprint density (RFP) in all 306 target mRNAs revealed that despite the reduction of footprints in the CDSes, translation was increased on the 5′ leaders in cells lacking DAP5 (Figure 1F), as measured by the ratio of footprints within the 5′ leader relative to the footprints at the annotated downstream CDS start codon. Increased translation in the 5′ leaders of the DAP5 target mRNAs occurred at uORFs as reflected by experimentally determined initiation-site-profiling in cells treated with harringtonine (HRT) and lactimidomycin (LTM) (Lee et al., 2012) (Figures 1E, G). In the presence of these drugs, ribosomes accumulate at the start codons but are allowed to complete elongation over the rest of the CDS (Ingolia et al., 2011; Lee et al., 2012). The majority of the DAP5 targets had multiple uORFs in the 5′ leader, with a median length of 26 codons, that frequently initiate at near-cognate start codons (CUG, GUG, UUG and AUC) in addition to the conventional AUG (Figures 1H-J). For instance, *WNK1*, a regulator of development and WNT signalling (Rodan and Jenny, 2017), exhibited increased ribosome occupancy in two GUG (one of which is in frame with the main CDS), two CUG, two UUG, one AUG and one ACG uORF (Figure 1E). These observations suggest that DAP5 mediates CDS, but not uORFs translation. Close inspection of the RFP profiles revealed that cap-proximal uORF translation is DAP5-independent (increased RFPs in null cells), whereas downstream uORFs and CDS are translated in a DAP5-dependent manner (decreased RFPs in null cells) (Figures 1E, F and S2).

Further analysis of the 5′ leader sequences of DAP5 target mRNAs also showed increased length, high GC-content and decreased minimum free energy (Figure S3A-C). In addition, DAP5-independent uORFs tend to concentrate in the regions of the 5′ leaders adjacent to high predicted propensity for structure (Figures 1E and S2). The increased complexity of the 5′ leader sequences of DAP5 target mRNAs was also associated with decreased TE of the main CDS (Figure S3D). These findings indicate that the 5′ leaders of DAP5 targets likely form structured elements that might define positional information for DAP5-dependent translation.

### Target mRNA 5′**′** leaders are necessary and sufficient to induce DAP5-dependent translation

Given that DAP5 acts in translation initiation (Liberman et al., 2015; Liberman et al., 2009; Marash and Kimchi, 2005; Sugiyama et al., 2017; Weingarten-Gabbay et al., 2014; Yoffe et al., 2016), we tested if the sequences of *WNK1*, *ROCK1* and *AKT1* 5′ leaders were sufficient to confer DAP5 sensitivity on a *Renilla* luciferase (R-LUC) reporter (Figure 2A-C). In comparison, the R-LUC luminescence driven by *WNK1*, *ROCK1* and *AKT1* 5′ leaders were reduced in DAP5-null cells to 20%, 30% and 40%, respectively (Figures 2A-C). Decreased translation of *WNK1-*, *ROCK1-* and *AKT1-*R-LUC reporters was not due to variations in mRNA abundance in the absence of DAP5 (Figures S3E-J). Re-expression of DAP5 (full length; FL) in the null cells restored R-LUC activity (Figures 2A-D, S3K-M), indicating that the 5′ leaders of *WNK1*, *ROCK1* and *AKT1* are sufficient to promote DAP5-dependent translation of R-LUC.

**Figure 2.**
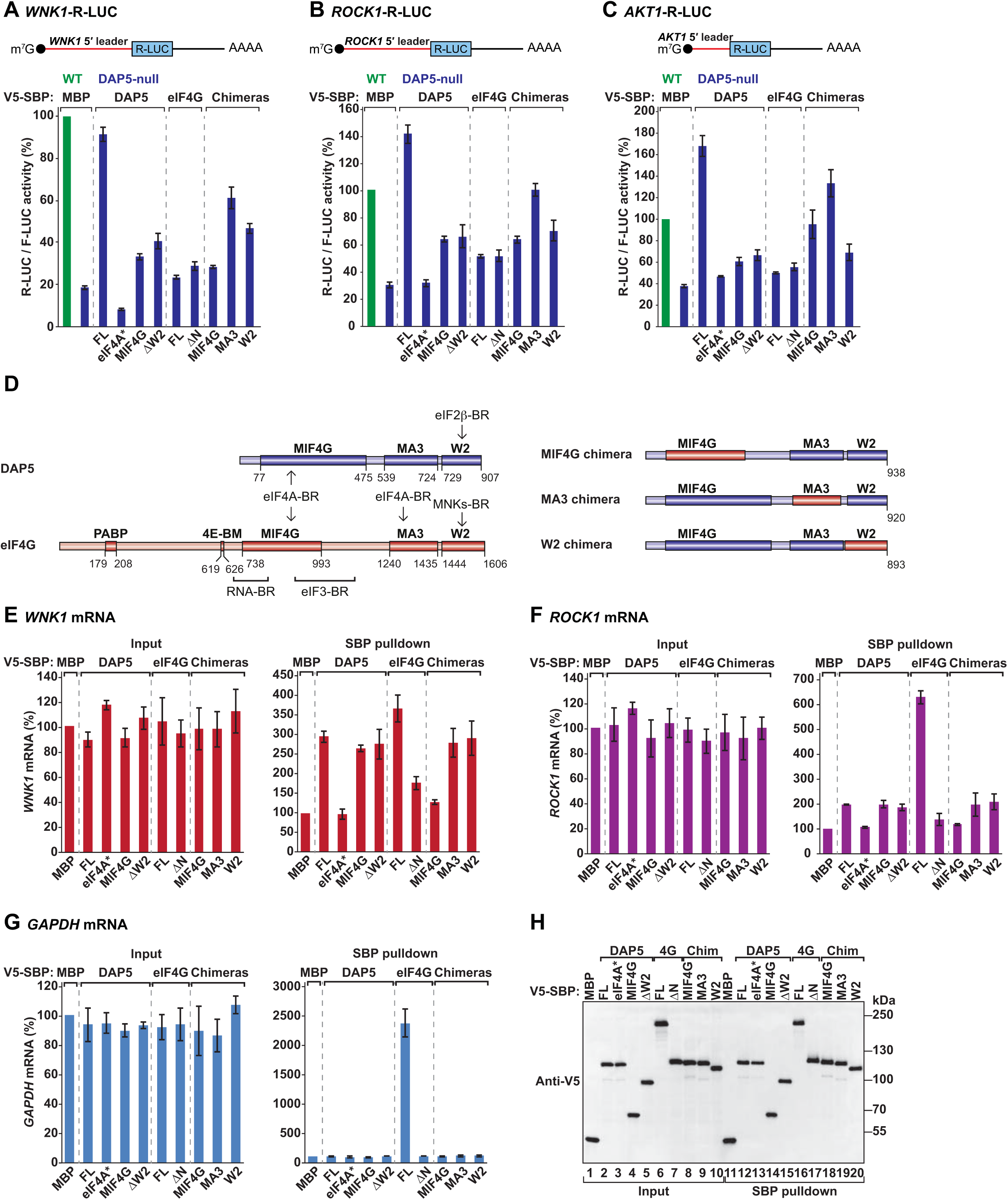
5′ leaders determine DAP5-dependent translation of target mRNAs. **(A-C)** WT (green) and DAP5-null (blue) HEK293T cells were transfected with different *Renilla* luciferase (R-LUC) reporters that contain the 5′ leader sequences of the *WNK1*, *ROCK1* and *AKT1* mRNAs cloned upstream of R-LUC CDS. Cells were also transfected with the normalization and transfection control F-LUC-GFP that contains a short 5′ leader sequence. In addition, the plasmids expressing V5-SBP tagged maltose binding protein (MBP), DAP5 [full length (FL) or the indicated mutants], eIF4G [full length (FL) or the indicated mutants], or DAP5-eIF4G chimeric proteins were also present in the transfection mixture. R-LUC activity was quantified in WT and DAP5-null cells in the presence of the different proteins two days post-transfection, normalized over to that of F-LUC-GFP and set to 100% in WT cells. The mean values +/- SD of three independent experiments are shown. Schematic representations of the R-LUC reporters are presented above each graph. Proteins are as follow: eIF4A*: eIF4A-binding mutant; MIF4G: DAP5 MIF4G domain; ΔW2: deletion of the DAP5 W2 domain; ΔN: deletion of eIF4G N-terminal region; MIF4G chimera: eIF4G MIF4G domain swapped into DAP5; MA3 chimera: eIF4G MA3 domain swapped into DAP5; W2 chimera: eIF4G W2 domain swapped into DAP5. See also Figure S3. **(D)** Schematic representation of the DAP5, eIF4G and DAP5-eIF4G chimeras. See Figure 1A for details. **(E-G)** HEK293T cells were transfected with plasmids expressing V5-SBP-tagged proteins: MBP, DAP5 (FL or the indicated mutants), eIF4G (FL or the indicated mutants), or DAP5-eIF4G chimeras. Streptavidin pulldown assays were performed two days post transfection and protein and RNA samples were obtained for each experimental condition. *WNK1*, *ROCK1* and *GAPDH* mRNA levels in input (0.8 %) and pulldown samples (12 %) were determined by quantitative PCR (qPCR) following reverse transcription. Values were set to 100% for V5-SBP-MBP. The mean values +/- SD of three independent experiments are shown. **(h)** Immunoblot depicting the expression and the pulldown efficiency of the V5-SBP tagged proteins used in E-G. Membranes were probed with anti-V5 antibody.

### DAP5-dependent translation requires the MIF4G and the W2 domains of the protein

To elucidate the molecular details of DAP5-dependent translation, we first tested if translation of the R-LUC reporters was influenced by changes in DAP5 protein sequence. We measured the activity of *WNK1-*, *ROCK1-* and *AKT1-*R-LUC in the null cells upon transient expression of DAP5 mutants. Translation was dependent on the interaction with the RNA helicase eIF4A, as R-LUC activity was not restored in the presence of DAP5 carrying mutations in the eIF4A-binding region (eIF4A*; Figures 2A-D, S4A). However, binding to eIF4A was not sufficient to induce DAP5-dependent translation, as the expression of the eIF4A-interacting domain (DAP5 MIF4G, residues 1-475) alone failed to restore R-LUC activity (Figures 2A-D, S3K-M, S4A). The interaction of DAP5 with eIF2β was also important for translation of the R-LUC reporters; R-LUC activity was still reduced in null cells expressing a DAP5 protein lacking the W2 domain (ΔW2, residues 1-722) and unable to associate with the β subunit of the ternary complex (Figures 2A-D, S3K-M, S4B).

We then asked if overexpression of full length (FL) or N-terminally truncated eIF4G (lacking the PABP and eIF4E binding sites; eIF4G ΔN) (Figure 2D) would suffice to translate the R-LUC reporters in the absence of DAP5. Curiously, none of the proteins was able to re-establish R-LUC activity (Figures 2A-D), indicating that *WNK1*, *ROCK1* and *AKT1* 5′ leaders drive translation of the main CDS in a DAP5-specific manner.

Lastly, we also used DAP5 chimeric proteins where the MIF4G, MA3 or W2 domains were swapped with the respective eIF4G domains (Figure 2D). Relative to the re-expression of DAP5 (FL), the chimeras were unable to fully restore R-LUC luminescence in the null cells, with the MIF4G and W2 chimeras showing the strongest defects on R-LUC translation (Figures 2A-D). These findings reinforce the notion that all domains of DAP5, and their specific interactions, are necessary for efficient translation of the R-LUC reporters with DAP5 target 5′ leaders. All DAP5 protein constructs were expressed at similar levels and reporter mRNA levels were not altered between the conditions (Figure S3E-M).

### Only eIF4A-bound DAP5 can interact with mRNA

To investigate the recruitment of DAP5 to target mRNAs, we performed RNA-pulldown assays and RT-qPCR. In contrast to V5-SBP-MBP, V5-SBP-DAP5 efficiently associated with *WNK1* and *ROCK1* mRNAs, but not *GAPDH* (Figure 2E-H). The association of DAP5 with the targets was abolished by mutations on the eIF4A-binding region (eIF4A*; Figure 2E, F), suggesting that eIF4A mediates mRNA binding. Consistent with this idea, the MIF4G domain of DAP5 was sufficient to pull down *WNK1* and *ROCK1* mRNAs, either alone (MIF4G) or when present in other DAP5 constructs (MA3 chimera and W2 chimera; Figure 2E-G). The DAP5 MIF4G was also specifically required for mRNA binding, as substitution by the respective domain in eIF4G, which is 39% sequence identical (Virgili et al., 2013), prevented DAP5 recruitment (Figure 2E-G). Consistent with a role of eIF4A in mRNA binding, we observed that one third of DAP5 targets (n=102; Figure S4C; Table S2) showed experimentally determined Rocaglamide A (RocA) sensitivity (Iwasaki et al., 2016). RocA is a translation inhibitor that clamps eIF4A onto polypurine sequences on the mRNA (Iwasaki et al., 2016). RocA-sensitive mRNAs, such as *WNK1*, show decreased RFP density at the CDS and premature uORF translation in the presence of the drug (Figure S4D, E) (Iwasaki et al., 2016). These findings indicate that DAP5 specifically interacts with transcripts when in complex with eIF4A. In the absence of an interaction with the RNA helicase, binding of DAP5 to the target, and thus translation, is compromised (Figures 2A-C).

The interaction of DAP5 ΔW2 with *WNK1* and *ROCK1* transcripts was comparable to the wild type protein (Figures 2E, F). This result suggests that impaired translation of the R-LUC reporters in null cells upon DAP5 ΔW2 expression (Figures 2A-C) is unrelated to target binding and is most likely associated with the function of the W2 domain in the initiation of translation.

In contrast to DAP5, eIF4G bound strongly to all tested mRNAs, including the DAP5 targets *WNK1* and *ROCK1*; however, its interaction with mRNA was dependent on its N-terminal region, which contains PABP-, eIF4E- and RNA-binding motifs (Figure 2D-G) (Imataka et al., 1998; Lamphear et al., 1995; Mader et al., 1995; Prevot et al., 2003; Yanagiya et al., 2009), as the N-terminally truncated eIF4G (ΔN) exhibited reduced mRNA binding. All proteins were expressed at equivalent levels and did not alter mRNA input levels (Figures 2E-G, input panels, H).

Altogether, our findings show that both eIF4G and DAP5 bind to *WNK1* and *ROCK1* mRNAs. Whereas the interaction of DAP5 with the mRNA is specific and reliant on eIF4A, eIF4G binds to all capped mRNAs as part of the eIF4F complex. Once on the mRNA, DAP5 mediates the synthesis of WNK1 and ROCK1 proteins (main CDS) but is dispensable for the translation of cap-proximal uORFs (Figures 1E, F). Translation of the latter is most likely eIF4G- and eIF4F-dependent. Thus, initiation of translation along the structured 5′ leaders of DAP5 targets switches from a DAP5-independent and eIF4F-dependent mechanism to a DAP5- and eIF4A-dependent mechanism.

### DAP5-mediated translation is cap-dependent

Given that the 5′ leaders of DAP5 targets contain structured elements that may represent IRESes promoting DAP5 recruitment, we blocked cap/eIF4F-dependent translation via the overexpression of an engineered eIF4E-binding protein (4E-BP) (Peter et al., 2015a) and tested binding of DAP5 to *WNK1*- and *ROCK1*-R-LUC mRNAs. As shown in cap-based pulldowns, overexpressed 4E-BP bound to eIF4E and abolished its interaction with eIF4G (Figure 3A), thus suppressing cap/eIF4F-dependent translation. Notably, binding of V5-SBP-DAP5 to the transcripts was suppressed in the presence of the 4E-BP (Figures 3B, C). The overexpressed proteins were pulled down at comparable levels in the different experimental conditions (Figure 3D). These findings suggest that binding of DAP5 to mRNA is cap/eIF4F-dependent and not mediated by an IRES-dependent mechanism.

**Figure 3.**
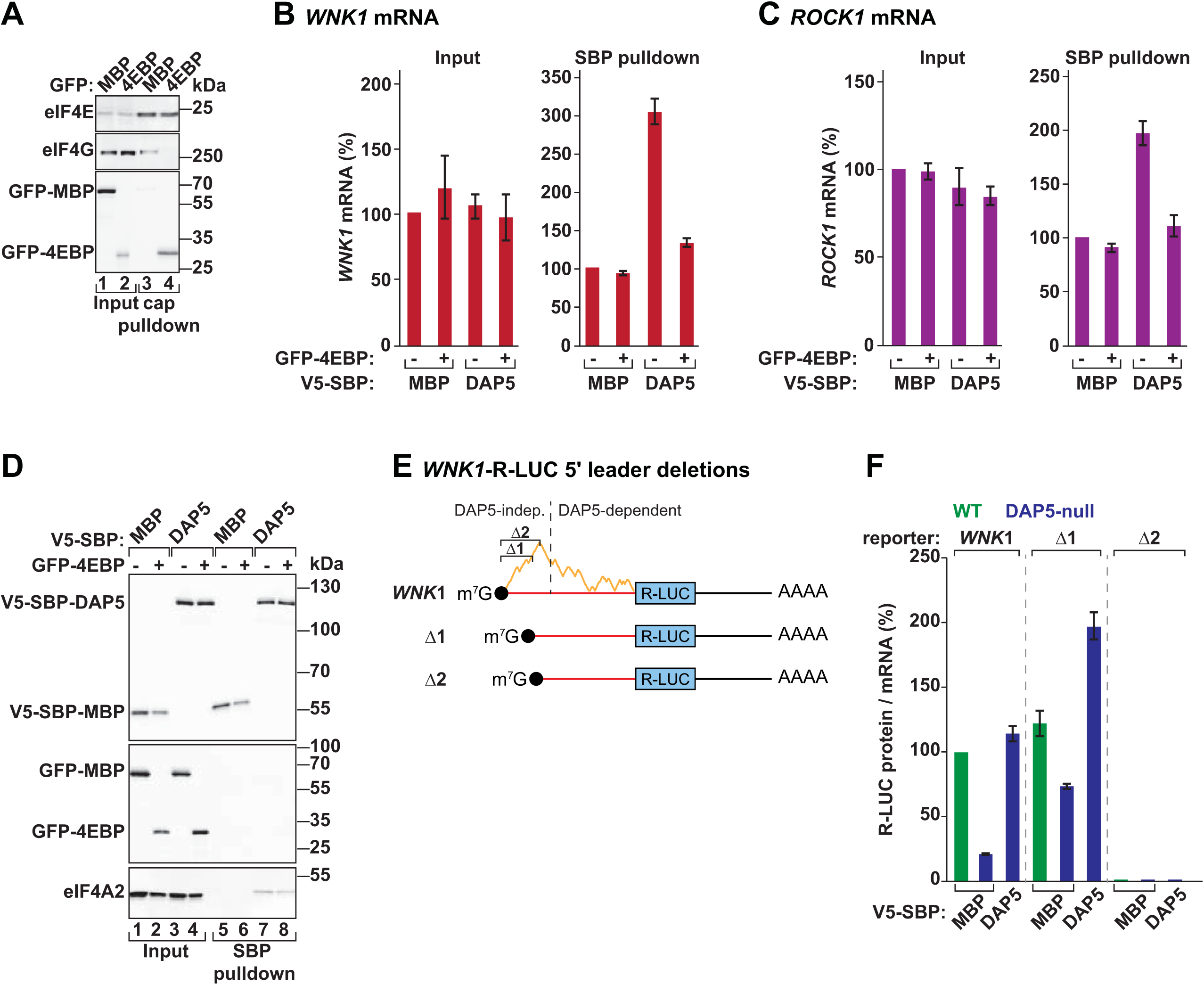
DAP5-dependent translation requires eIF4F-mediated ribosome recruitment. **(A)** m^7^GTP-cap pulldown assay showing the interaction between eIF4E and eIF4G in the presence or absence of GFP-4EBP. Inputs (1% for eIF4E and 0.3% for eIF4G and GFP-tagged proteins) and bound fractions (1% for eIF4E and 2% for eIF4G and the GFP-tagged proteins) were analysed by western blotting. Membranes were probed with anti-eIF4E, eIF4G and GFP antibodies. **(B, C)** Binding of V5-SBP-DAP5, or V5-SBP-MBP as control, to *WNK1* and *ROCK1* mRNAs was determined by RNA-immunoprecipitation (IP) in the presence or absence of GFP-4EBP. Proteins were pulled down using streptavidin beads. mRNA levels in input (0.8%) and IP samples (12%) were quantified by RT-qPCR and set to 100% for V5-SBP-MBP. Bars indicate the mean value; error bars represent SD (n=3). **(D)** Immunoblot depicting the expression of the proteins used in the RNA-IP. Inputs were 1% for the V5-SBP-tagged proteins and 0.3% for the GFP-tagged and endogenous proteins. Bound fractions correspond to 1% for the V5-SBP-tagged proteins and 2% for the GFP-tagged and endogenous proteins. eIF4A2 was used as a positive interactor of DAP5. Blots were probed with anti-V5, GFP and eIF4A2 antibodies. **(E)** Schematic representations of the *WNK1*-R-LUC reporters with cap-proximal deletions that that partially (Δ1) or completely (Δ2) remove the uORFs translated independently of DAP5 and located upstream of the structure-prone region of the 5′ leader. The predicted propensity for secondary structure across *WNK1* 5′ leader is illustrated in orange. **(F)** WT and DAP5-null cells were transfected with the *WNK1-*R-LUC reporters represented in (E), F-LUC-GFP, and V5-SBP-MBP or V5-SBP-DAP5. Following transfection, luciferase activities (Protein) were measured and mRNA levels determined by northern blotting. R-LUC values were normalized to the transfection control F-LUC-GFP. The graph shows the protein to mRNA ratios in WT and null cells, set to 100% in WT cells expressing the *WNK1-*R-LUC reporter. See also Figure S4.

We then generated two cap-proximal truncations in *WNK1-*R-LUC mRNA (Δ1 and Δ2; Figure 3E). These truncations partially (Δ1) or completely (Δ2) removed the sequence in the 5′ leader containing the cap-proximal uORFs translated independently of DAP5. Both truncations reduced the abundance of the reporter, and consequently R-LUC activity (Figure S4F-H), suggesting they might affect mRNA stability and/or transcription. To assess changes only in translation, we determined the protein/mRNA ratios (TE) for the *WNK1-*R-LUC Δ1 and Δ2 reporters. We observed that despite low mRNA levels, the Δ1 reporter was still translated with the same efficiency, even in the absence DAP5 (Figure 3F and S4I). *WNK1-*R-LUC Δ2 mRNA was not translated. These findings show that the cap-proximal region of the 5′ leader is critical for DAP5-mediated translation. This result also indicates that the downstream structured region is not sufficient to determine the translation of the main CDS. Since cap-proximal uORFs are translated in the null cells (Figure 1D), we conclude that DAP5 acts downstream of cap-dependent translation initiation, i.e., DAP5 drives translation of the main CDS in mRNAs where eIF4F-loaded ribosomes translate uORFs.

### DAP5 enables re-initiation after uORF translation

Our data suggests that the uORFs located in the 5′ leaders of DAP5 targets serve an important role in the translation of the main CDS by DAP5. To characterize the functionality of uORFs in the regulation of DAP5-dependent translation, we transfected wild type and null cells with versions of the *WNK1-*R-LUC reporter containing altered uORF features.

The *WNK1* transcript contains at least eight uORFs, one of which (uORF2) is in frame with the AUG of the main CDS. uORF2 is translated in the absence of DAP5, initiates from a GUG start codon (uGUG) and is 22 codons in length (Figures 1E and 4A). We optimized the initiation sequence context of uORF2 by mutating the uGUG to conventional AUG (uORF2+; Figure 4A). Based on the proximity to the 5′ end, the uAUG should play a dominant role in start codon recognition and in the initiation of translation (Kozak, 2002). Interestingly, *WNK1*-R-LUC-uORF2+ mimicked the reporter with the natural 5′ leader of *WNK1* (*WNK1*): it promoted the expression of R-LUC (main CDS) in a DAP5-dependent manner (Figures 4B, C, lanes 1-6). This result suggested that uORF2 translation sustained the expression of the main CDS.

**Figure 4.**
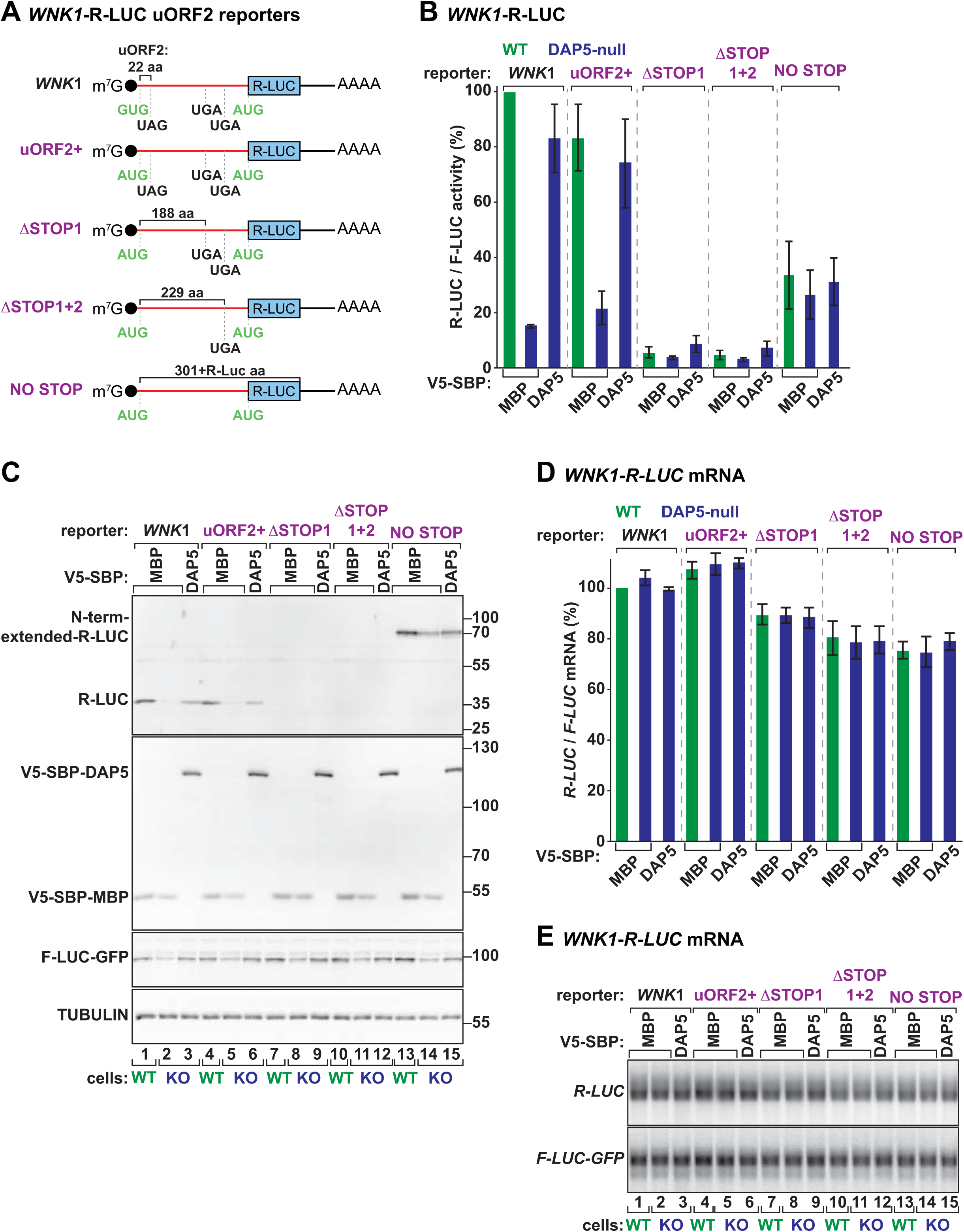
DAP5 mediates re-initiation following uORF translation. **(A)** Schematic representations of the *WNK1*-R-LUC reporters with changes in uORF2 initiation context and length. uORF2 GUG is in frame with the R-LUC AUG. uORF2 is 22 codons long. Three STOP codons can be found downstream and in frame with uGUG. uORF2+: GUG start codon was substituted by AUG to favour initiation of translation. ΔSTOP1: first STOP codon in frame with uAUG is removed; uORF2 is then 188 codons long. ΔSTOP1+2: reporter lacks the two STOP codons after the uAUG; length of uORF2 increases to 229 codons. NO STOP: the three STOP codons are absent; the reporter produces a N-terminally extended R-LUC. aa: amino acids. **(B-E)** WT and DAP5-null cells were transfected with the different *WNK1*-R-LUC reporters shown in A, F-LUC-GFP and V5-SBP-MBP or V5-SBP-DAP5. After transfection, R-LUC activity was measured, normalized to F-LUC-GFP and set to 100% for WT cells expressing *WNK1*-R-LUC (B). Expression of short and long (N-terminally extended) R-LUC was also evaluated by immunoblotting, together with V5-tagged proteins, F-LUC-GFP and TUBULIN using anti-R-LUC, V5, GFP and TUBULIN antibodies (C). The abundance of the different *R-LUC* and *F-LUC* reporter mRNAs was assessed by northern blotting and quantified as in B (D, E). Bars indicate the mean value; error bars represent SD (n=3). See also Figure S5.

To confirm that the uAUG was used in the initiation of translation, we removed all STOP codons, and consequently all uORFs, in frame with the AUG of R-LUC (3 in total) (NO STOP; Figure 4A). In the *WNK1*-R-LUC-NO STOP reporter, the uAUG was the only initiating codon and originated an R-LUC protein with an extended N-terminal region (70 kDa instead of 35 kDa; Figure 4C, lanes 13-15 vs 1-3). Although the protein levels of the two forms of the luciferase were similar (Figure 4C), the N-terminal extension reduced R-LUC activity (Figure 4B). Moreover, the activity of the long R-LUC was similar in wild type and null cells, indicating that its translation was not mediated by DAP5 (Figures 4B).

We also generated reporters where only one (ΔSTOP1) or two (ΔSTOP1+2) of the three STOP codons were removed. Single and double deletions of these STOPs increase the size of uORF2 to 188 or 229 codons, respectively (Figure 4A). In these settings R-LUC translation was abolished (Figures 4B, C), indicating that DAP5-dependent translation of the main CDS is influenced by uORF length, as only the short uORF2 (22 codons in length) permitted R-LUC translation. To test the maximum uORF2 length supporting DAP5-mediated translation of R-LUC, we extended the position of its STOP to 29, 39 or 49 codons downstream of uAUG (Figure S5A). Although all tested reporters sustained DAP5-dependent translation of R-LUC, the expression and activity of the luciferase was inversely correlated with uORF2 length (Figures S5B, C). None of the observed differences could be explained by varying mRNA levels (Figures 4D, E and S5D, E). The fact that R-LUC synthesis is only observed in the presence of short uORFs shows that DAP5 drives re-initiation of translation.

We also examined the ability of DAP5 to prime re-initiation of translation without altering uORF initiation context. In the *WNK1* 5′ leader, uORF6 initiates with a conventional AUG (uAUG), is translated in a DAP5-dependent manner and is in frame with a UGA STOP located 4 codons downstream (Figures 1E, 5A). uORF6 translation occurs in the –1 frame. We interfered with termination at the UGA codon and produced reporters encoding uORF6 with distinct lengths: 118, 30, 19 or 9 codons (Figure 5A). The *WNK1* reporters with the engineered 5′ leaders were transfected into cells and assayed for R-LUC activity and expression (Figures 5B, C and S5F-H). As observed above, DAP5-dependent translation of *R-LUC* was regulated by uORF length. In the presence of a long uORF6 (uORF118 and uORF30) or uORF19, *R-LUC* was weakly translated (Figures 5B, C, lanes 4-12 vs 1-3). Short uORF6 length (uORF9) primed *R-LUC* translation in a DAP5-dependent manner although with less efficiency than the natural uORF6 which encodes a 4 amino acid peptide (Figures 5B, C, lanes 10-15). Changes in mRNA abundance of the different reporters were not sufficient to explain the variation in the efficiency of R-LUC translation (Figures 5B, S5F-H). These findings are consistent with a model where DAP5 promotes translation of main CDSes after short uORF translation.

**Figure 5.**
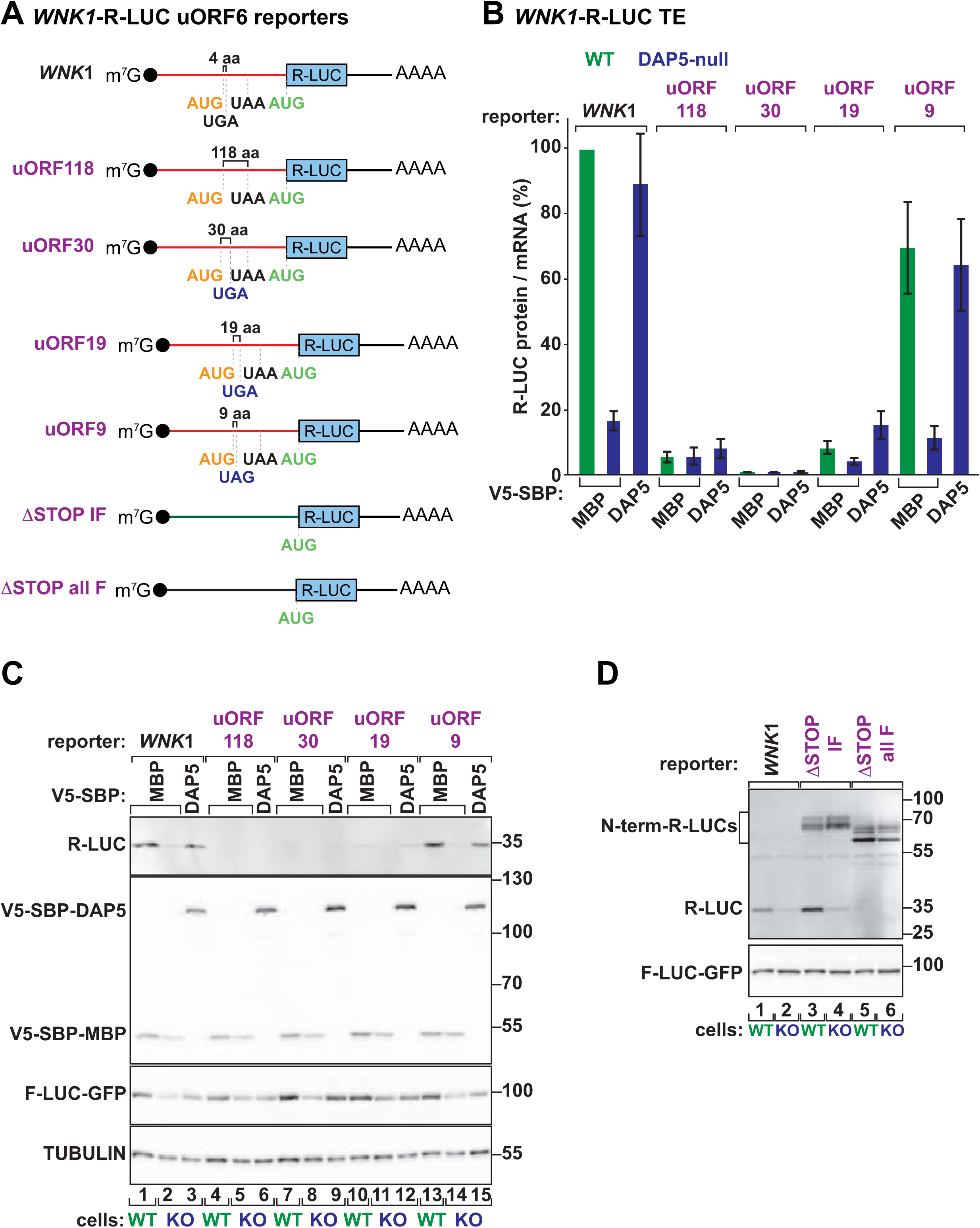
uORF length is critical for re-initiation of translation by DAP5. **(A)** Schematic representations of the WNK1-R-LUC reporters with changes in uORF6 length. uORF6 initiates from an AUG start codon in the –1 frame and encodes a short peptide (4 amino acids). uORF118: UGA STOP codon was removed changing the length of uORF6 to 118 codons. uORF30, uORF19, uORF9: position of the STOP codon was moved to 30, 19 or 9 codons downstream of uAUG, respectively. ΔSTOP IF (in-frame): all STOP codons in frame with uAUG were removed. ΔSTOP all F (frames): the *WNK1* 5′ leader lacks STOP codons. **(B, C)** WT and DAP5-null cells were transfected with different *WNK1*-R-LUC reporters, F-LUC-GFP and V5-SBP-MBP or V5-SBP-DAP5. Following transfection, luciferase activities (Protein) were measured and mRNA levels determined by northern blotting. R-LUC values were normalized to the transfection control F-LUC-GFP. The graph shows the protein and mRNA ratios in WT and null cells, set to 100% in WT cells expressing *WNK1-*R-LUC. See also Figure S5. The immunoblot showing the expression of the different proteins is shown in panel C. TUBULIN served as a loading control. **(D)** Western blot depicting the expression of short and long (N-terminally extended) R-LUC proteins produced in WT and DAP5-null cells transfected with the *WNK1*-R-LUC ΔSTOP IF and *WNK1*-R-LUC ΔSTOP all F plasmids. F-LUC-GFP served as a transfection control.

Lastly, to exclude the possibility that DAP5 promotes translation initiation on downstream CDSes using PICs that scan past the uORFs, we determined the changes in R-LUC translation if the natural 5′ leader lacks all STOP codons in frame with the main CDS (ΔSTOP IF; Figure 5A). When transfected into cells, this reporter originated R-LUC proteins with different sizes (with and without distinct N-terminus), as observed by immunoblotting (Figure 5D, lane 3). The synthesis of several N-terminally extended versions of R-LUC indicates that the PICs scanning the 5′ leader initiate translation at different upstream start codons, as expected by leaky scanning at near-cognate start codons. In the null cells though, expression of short R-LUC (35 kDa), but not the majority of the long R-LUCs with N-terminal extensions, was diminished (Figure 5D, lane 4). We also removed all STOPs from the 5′ leader of *WNK1* (ΔSTOP all F; Figure 5A). In this reporter devoid of uORFs, translation can initiate at multiple start codons, but does not terminate before the main CDS encoding R-LUC. In cells, long R-LUC proteins were expressed independently of DAP5 (Figure 5D, lanes 5 and 6). Even if engineering of *WNK1* 5′ leader sequence was performed without introducing new start codons or altering the initiation contexts of the different uORFs (see methods section for details), removal of STOP codons changed start codon recognition, as shown by the presence of long R-LUC proteins with distinct sizes (Figure 5D, lanes 5,6 vs 3,4). Notably, short R-LUC was not synthesized in wild type and null cells.

These results have several implications. First, translation of the main CDS (R-LUC) in the context of *WNK1* 5′ leader is DAP5-dependent. Second, initiation at the main AUG only occurs after uORF translation. Third, DAP5 is critical for re-initiation of translation at the main CDS. Lastly, in the *WNK1* 5′ leader the PICs are unable to scan until the main AUG, signifying that DAP5 reutilizes the ribosomal complexes involved in uORF translation.

### Simultaneous uORF and main CDS translation in the DAP5 targets

The luciferase-based reporters used in the previous experiments suggest that uORF translation is pervasive and necessary for DAP5-dependent translation of the main CDS. However, in these experiments we are unable to detect the synthesis of uORF-derived peptides, and therefore confirm uORF translation. To simultaneously detect and quantify uORF and main CDS translation, we adopted a split-fluorescent protein approach using mNeonGreen2 (mNG2) that expresses the yellow-green-coloured protein in two fragments: mNG2_1-10_ and mNG2_11_. mNG2_1-10_ originates a non-fluorescent mNG2 due to the lack of 11^th^ β-strand; however, upon co-expression with mNG2_11_ (16-aa peptide), the two fragments assemble a functional mNG2 molecule (Chen et al., 2020; Feng et al., 2017; Leonetti et al., 2016). The uORF2 (22 aa) in the *WNK1* 5′ leader was replaced with the mNG2_11_ CDS initiating with an uAUG. In addition, the main CDS encoded the EBFP (enhanced blue fluorescent protein) (Figure 6A). The split-fluorescent reporters were transfected into wild type and DAP5-null cells together with a transfection control expressing mCherry.

**Figure 6.**
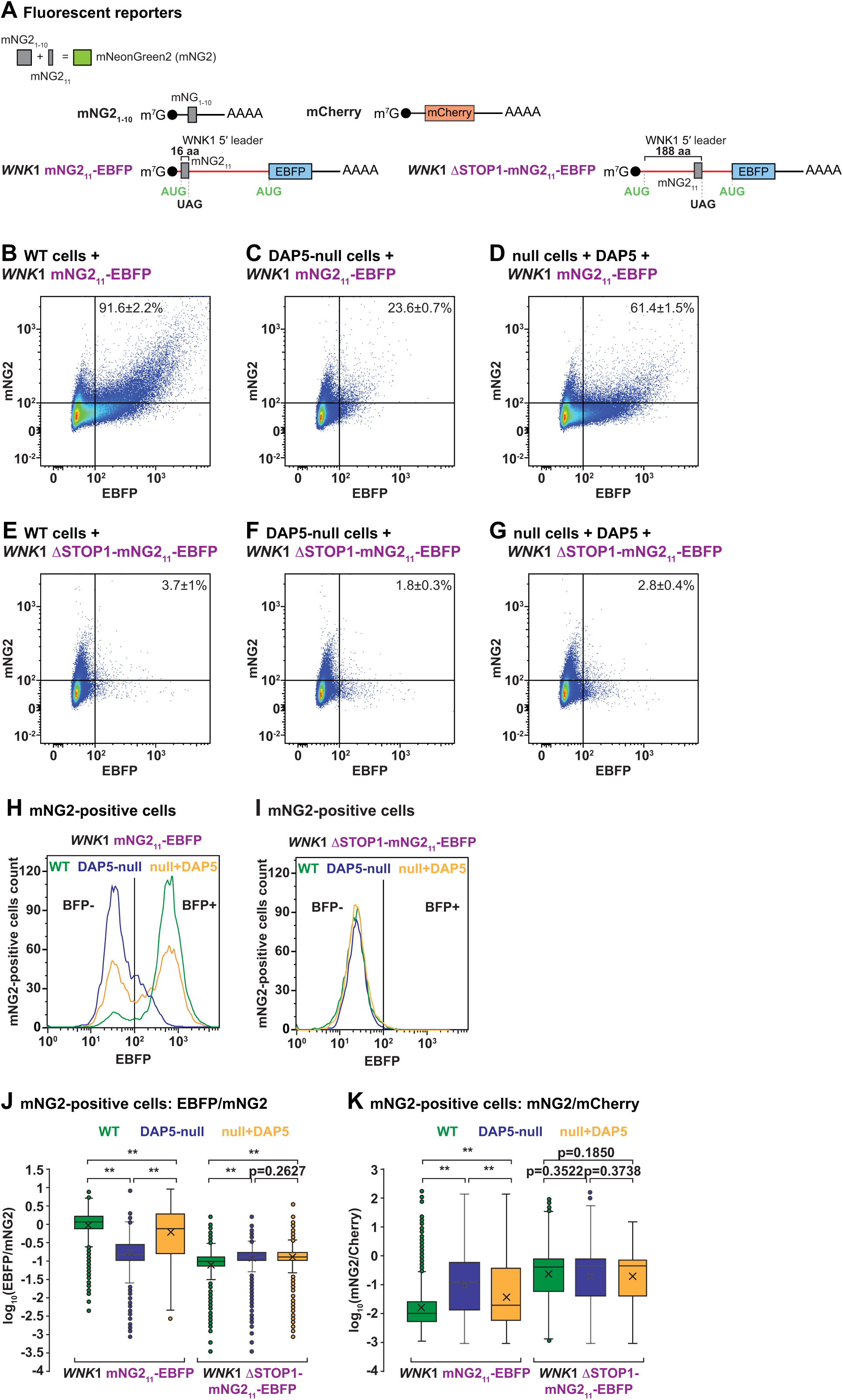
Concurrent uORF and main CDS translation in DAP5 targets. **(A)** Schematic representation of the mNeonGreen2 (mNG2) split-fluorescent protein approach and corresponding reporter constructs. Co-expression of the mNG2_1-10_ and mNG2_11_ fragments originates a functional mNG2 fluorescent molecule (Chen et al., 2020; Feng et al., 2017; Leonetti et al., 2016). mNG2_11_ CDS was inserted in *WNK1* 5′ leader and replaced uORF2. mNG2_11_ translation initiates at a uAUG in frame with the main CDS and produces a 16 aa protein. Main CDS encoded the EBFP fluorophore. ΔSTOP1: The first UAG STOP codon after the uAUG was removed and the mNG2_11_ CDS was inserted next to the UAG STOP located 188 codons downstream of the uAUG. **(B-I)** WT and DAP5-null cells were transfected with the mNG2_1-10_, *WNK1*-mNG2_11_-EBFP or *WNK1* ΔSTOP1-mNG2_11_-EBFP, mCherry, and V5-SBP-MBP or V5-SBP-DAP5 plasmids. Following transfection, cells were collected and analysed by flow cytometry. The scattered plots in panels b to g show the EBFP and mNG2 signal intensity in all measured cells in the presence (WT cells, null cells+DAP5) or absence (DAP5-null cells) of DAP5. mNG2 and EBFP expression are plotted on a bi-exponential scale and represent around 160000 cells. The values in the panels represent the proportion of mNG2-positive cells that were also EBFP-positive. In panels H and I, the histograms show the EBFP signal intensity in mNG2-positive cells (5000 cells in H and 3700 cells in I) detected in experiments B to G. EBFP expression is plotted on a log scale. **(J, K)** Box plots of the EBFP/mNG2 and mNG2/mCherry ratios quantified by flow cytometry in WT or DAP5-null cells, and null cells following V5-SBP-DAP5 re-expression. Cells expressed the mNG21-10, WNK1-mNG211-EBFP or WNK1 ΔSTOP1-mNG211-EBFP, and mCherry reporters. Boxes represent the 25th to 75th percentiles; black line shows the median and the cross the average; whiskers show the variability outside the upper and lower quartiles; dots show the outliers. Significance determined by one-sided Wilcoxon rank-sum test and indicated if p<2.2e-16 (**). null+DAP5: DAP5-null cells re-expressing V5-SBP-DAP5. See also Figure S6.

The non-overlapping excitation and emission spectra of the three fluorophores allowed their simultaneous detection by flow cytometry (Figure S6A-D). Expression of the mNG2 plasmids in *trans* did not generate a yellow-green signal (Figure S6A). Co-expression of the two mNG2 plasmids generated the fluorescent signal in up to 9% of the cells (Figure S6B). Although the complementation efficiency of the split-mNG2 system was low compared to the transfection efficiency in HEK293T cells (∼50% in wild type cells and ∼36% in DAP5- null cells, as assessed by the number of mCherry-positive cells, Figure S6D) it clearly showed that uORF translation (mNG2-positive) occurs in the *WNK1* 5′ leader (Figures 6B and S6B). EBFP fluorescence was only detected in cells expressing the *WNK1* mNG2_11_+BFP reporter (Figure S6C).

Close inspection of the fluorescent output in cells expressing the two mNG2 plasmids showed that the majority of mNG2-positive cells were also EBFP-positive (Figure 6B), indicating that uORF2 and main CDS are simultaneously expressed. The *WNK1* mNG2_11_+EBFP reporter also recapitulated DAP5-dependent translation of the main CDS. In the absence of DAP5, a large portion of cells expressing a functional mNG2 (uORF2) do not express EBFP (main CDS) (Figures 6C, H, J). The number of mNG2 and EBFP double positive cells was restored upon re-expression of DAP5 in the null cells (Figures 6D, H, J). Consistent with a block in re-initiation following translation of long uORFs, EBFP fluorescence was reduced and DAP5-independent in cells expressing the *WNK1* ΔSTOP1+mNG2_11_+EBFP reporter which encodes a mNG2_11_ peptide fused to 188 amino acids (Figures 6E-G, I, J). mNG2 expression (uORF translation) did not require DAP5 and was not disturbed by uORF length (Figure 6K). These observations confirm that short uORF translation in the 5′ leader of DAP5 targets promotes main CDS translation. Another implication of our results using different reporter systems is that uORF2 or main CDS sequences and peptides are not relevant for the re-initiation of translation by DAP5, excluding the possibility that uORF translated peptides influence CDS expression in *cis*. These experiments do not dismiss however, that *WNK1* uORFs-derived peptides are functional in cells.

### DAP5 utilizes post-termination translation complexes

To further test translation re-initiation by DAP5, we interfered with termination by exploiting a dominant negative mutant of the release factor 1, eRF1^AAQ^ (Brown et al., 2015; Shao et al., 2016), to cause local translation arrest at STOP codons. eRF1^AAQ^ is unable to hydrolyse the peptidyl-tRNA after STOP codon recognition (Frolova et al., 1999). Cells were transfected with the *WNK1*-R-LUC and GFP-F-LUC in the absence or presence of increasing amounts of eRF1^AAQ^ and luciferase activities and expression were measured. As expected upon termination inhibition, eRF1^AAQ^ expression decreased R-LUC and GFP-F-LUC protein levels in a concentration-dependent manner (Figure 7B, lanes 1-4). However, the R-LUC: F-LUC activity ratio varied if R-LUC translation was primed or not by DAP5. In the context of *WNK1* 5′ leader (*WNK1*-R-LUC and *WNK1*-R-LUC-uORF2+), increasing levels of eRF1 proportionally decreased R-LUC activity (Figures 7A-C). In contrast, DAP5-independent translation of R-LUC using a reporter containing a short 5′ leader (R-LUC) or an engineered *WNK1* 5′ leader without STOP codons that leads to the synthesis of the N-terminally extended R-LUC (*WNK1-*NO STOP-R-LUC, Figure 7A), was less affected by the eRF1^AAQ^ mutant. In these cases, R-LUC: F-LUC ratios were constant or even increased in the presence of rising levels of the mutant release factor (Figures 7B, C). In all the conditions, R-LUC mRNA levels remained unchanged (Figures S6E, F). These observations suggest that inhibition of termination after uORF translation impairs DAP5-dependent re-initiation at the main CDS.

**Figure 7.**
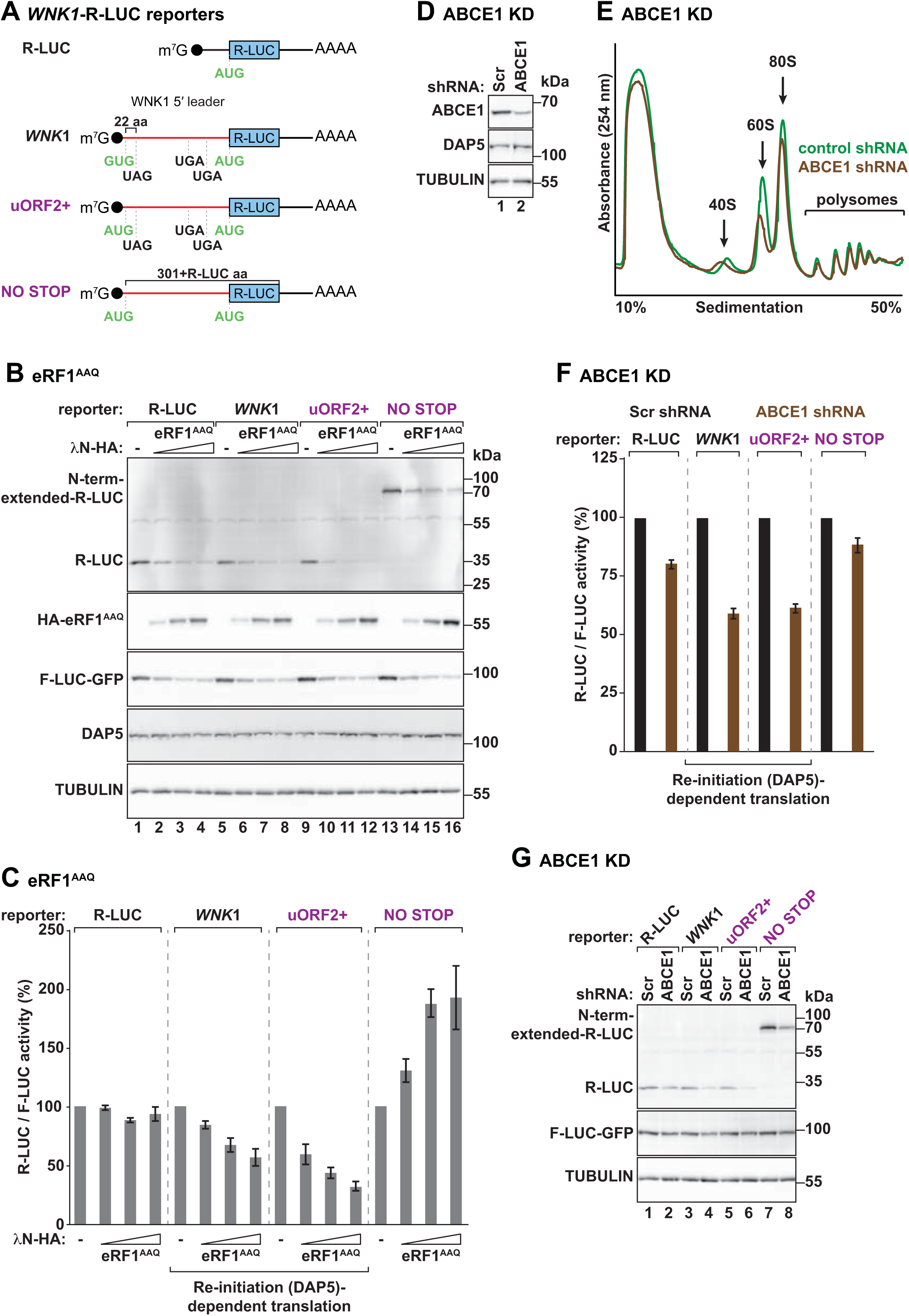
DAP5 uses post-termination translation complexes. **(A)** Schematic representation of *WNK1*-R-LUC reporters with changes in uORF2 length, as described in Figure 4. **(B, C)** HEK293T cells were transfected with the *WNK1-*R-LUC reporters shown in A. Additionally, the transfection mixtures also contained F-LUC-GFP and increasing concentrations of λN-HA-eRF1 . (B) Immunoblot showing the levels of the expressed proteins. The membranes were blotted with anti-R-LUC, HA, GFP, DAP5 and TUBULIN antibodies. (C) R-LUC activity was measured, normalized to F-LUC-GFP and set to 100% in the absence of λN-HA-eRF1 for each reporter. Bars indicate the mean value; error bars represent SD (n=3). See also Figure S6. **(D)** Western blot showing shRNA-mediated depletion of ABCE1 in HEK293T cells. TUBULIN served as a loading control. DAP5 expression did not vary in the absence of ABCE1. **(E)** UV absorbance profile at 254 nM of scramble shRNA (control) and ABCE1-depleted (ABCE1 shRNA) HEK293T cell extracts after polysome sedimentation in a sucrose gradient. 40S and 60S subunits, 80S monosomes, and polysome peaks are indicated. **(F, G)** HEK293T cells were treated with scramble (Scr) or shRNA targeting *ABCE1* mRNA and transfected with the *WNK1*-R-LUC reporters shown in A. (F) The graph shows relative R-LUC activity in control (Scr) and ABCE1 KD cells. R-LUC activity was normalized to that of F-LUC-GFP and set to 100% in Scr-treated cells for each reporter. (G) Immunoblot illustrating the expression of short and long (N-terminally extended) R-LUC, F-LUC-GFP and TUBULIN in control and ABCE1-depleted cells. Blots were probed with anti-R-LUC, GFP and TUBULIN antibodies. See also Figure S6.

In agreement with the re-initiation model, similar findings were obtained when 60S recycling was impaired in cells expressing the *WNK1*-R-LUC reporters. As expected, shRNA-mediated depletion of the ATP binding cassette sub-family E member 1 (ABCE1 KD; Figure 7D) decreased the levels of free 60S subunits in cells, as judged in polysome profiles of control (scramble) or ABCE1 shRNA-treated cells after sucrose density gradient separation (Figure 7E). In cells with low levels of ABCE1, DAP5-dependent re-initiation of R-LUC translation (*WNK1*-R-LUC and *WNK1*-uORF2+-R-LUC reporters) was pronouncedly decreased compared to DAP5-independent translation of R-LUC (R-LUC and *WNK1*-NO STOP-R-LUC reporters) (Figures 7F, G). Depletion of ABCE1 did not affect the levels of the different R-LUC transcripts (Figures S6G, H). These findings indicate that DAP5 acts on post-termination translation complexes following 40S and 60S subunits dissociation at the STOP codons of uORFs.

### DAP5 targets partially overlap with re-initiation dependent transcripts

eIF2D and the Density-regulated protein (DENR)/Malignant T-cell amplified sequence-1 (MCTS-1) complex recycle 40S ribosomal subunits (Bohlen et al., 2020; Young et al., 2018). In animals, these non-canonical translational factors also promote re-initiation (Ahmed et al., 2018; Bohlen et al., 2020; Castelo-Szekely et al., 2019; Schleich et al., 2017; Schleich et al., 2014; Vasudevan et al., 2020). The underlying mechanism for the function of these proteins in translation re-initiation remains controversial as different functions have been described for these proteins. eIF2D, DENR and MCTS-1 have been proposed to act as non-canonical Met-tRNA_i_ delivery factors (Skabkin et al., 2010; Vasudevan et al., 2020) or as translation factors promoting the eviction of deacylated tRNAs that interact strongly with the post-termination 40S complexes on STOP codons (Bohlen et al., 2020; Skabkin et al., 2010).

To exploit if altered function of these factors affects re-initiation of translation by DAP5, we depleted the proteins in cells using shRNA-mediated KD (Figures S7A). Cells were then transfected with the reporters driving R-LUC (main CDS) synthesis by re-initiation (DAP5)-dependent and independent mechanisms (Figure S7B). In the absence of DENR, MCTS-1 and eIF2D, R-LUC expression from all reporters was slightly reduced (Figures S7C, D), suggesting that these factors are not specifically required for re-initiation of translation by DAP5 in the context of *WNK1* 5′ leader. However, as DENR/MCTS-1 and eIF2D have been shown to selectively support re-initiation after certain uORFs (Ahmed et al., 2018; Bohlen et al., 2020; Castelo-Szekely et al., 2019; Schleich et al., 2017; Schleich et al., 2014), we compared the DAP5 targets with the group of mRNAs showing reduced translation in DENR knockout HeLa cells (Bohlen et al., 2020). Approximately 20% of the DAP5 targets (excluding *WNK1*) were dependent on DENR for efficient translation (Figure S7E, Table S3). These results suggest that re-initiation of translation by DAP5 requires DENR activity following uORF translation in a specific group of mRNAs. As DENR/MCTS-1 and eIF2D are either non-canonical Met-tRNA_i_^Met^ delivery factors or deacylated tRNA eviction factors (Bohlen et al., 2020; Skabkin et al., 2010; Vasudevan et al., 2020), this data also implies that re-initiation of translation by DAP5 utilizes deacylated tRNA-free 40S subunits bound to the mRNA.

Altogether, our work shows that DAP5 mediates re-initiation following uORF translation. The data support a model in which binding of the DAP5-eIF4A complex to 40S subunits following termination of uORF translation promotes re-initiation at the downstream CDS. The structured nature and the presence of multiple uORFs in the 5′ leaders of DAP5 targets may favour dissociation of the eIF4F complex. In this context, translation of cap-distal CDSes relies on DAP5. In the absence of this non-canonical eIF4G protein, post-termination 40S complexes are most likely prone to dissociate from intricate 5′ leaders.

## Discussion

Here we reveal that DAP5 is a non-canonical factor that mediates re-initiation without affecting general cap-dependent translation. As one of the few re-initiation specific factors described to date, DAP5 emerges as an important protein in translational control with multiple biological implications. Re-initiation-dependent transcripts are enriched for regulatory proteins such as kinases and phosphatases, implicating DAP5 in the control of cell signalling cascades that support cell proliferation and differentiation. Our data, also expands the list of mRNAs in which re-initiation of translation is essential for protein synthesis.

### DAP5 enhances re-initiation on mRNAs with burdened 5′ leaders

The cues for DAP5-mediated translational control are driven by information present in the 5′ leaders of its target mRNAs. Transcripts with structure-prone 5′ leaders and multiple uORFs selectively require DAP5 for proper translation of the main CDS. The long and burdened 5′ leaders restrain scanning of cap-loaded pre-initiation complexes, facilitate translation at uORFs that otherwise would be skipped, and limit main CDS translation (Kozak, 1990). Re-utilization of post-termination complexes following uORF translation is therefore instrumental to trigger the synthesis of the proteins encoded by the main CDS. In this scenario, DAP5 plays a unique role: together with eIF4A, it binds to the mRNA and enables translation re-initiation. We propose that repeated uORF translation and scanning cycles fuelled by DAP5 at the impenetrable 5′ leaders move the ribosome towards the main CDS (Figure S7F model).

Mechanistically, DAP5 most likely replaces the function of the eIF4F complex. The intricate nature of scanning coupled with slow translation of sequenced biased (GC-rich) uORFs might gradually dissociate or reduce the activity of eIF4F along the long 5′ leaders and favour binding of DAP5 to 40S subunits. As DAP5 interacts with eIF4A, eIF3 and eIF2β (Imataka et al., 1997; Lee and McCormick, 2006; Liberman et al., 2015), its presence on the mRNA may stabilise post-termination 40S subunits, promote the recruitment of an initiator tRNA and/or stimulate a new cycle of scanning and translation. Indeed, DAP5 mutant proteins unable to associate with these initiation factors exhibited reduced ability to promote re-initiation, and eIF4G or its N-terminally truncated protein did not substitute DAP5 in null cells. Future studies will enable the detailed characterization of DAP5 functions as a re-initiation factor.

We present additional evidence that supports the proposed model for the role of DAP5 in translational control. Loss of DAP5 only affected the translational efficiency of a fraction of the transcriptome. Ribosome footprint profiles of DAP5 targets and target-based luciferase reporters indicated prevalent uORF, but not main CDS translation in the null cells. Increased ribosome density on the uORFs at the expense of the main CDS is consistent with an inability of ribosomes to re-initiate downstream of uORF translation. Failure to re-initiate in the absence of DAP5 can be associated with impediments to scanning by the stable secondary structures on the 5′ leader, and the absence of cofactors that license start codon recognition and 80S formation, or stabilize the interaction of post-termination complexes with the mRNA. Accordingly, polysome profiling assays in DAP5-null cells show increased levels of free 40S subunits.

DAP5 recruitment to the mRNA was determined by the 5′ leader sequence, ribosome loading, and binding to eIF4A. Thus, DAP5 acts upon the initiation of cap-dependent translation on mRNAs that depend strongly on eIF4A for scanning. Understanding the dynamics of eIF4F and DAP5 association/dissociation with the translation machinery will highlight the interplay of different initiation complexes in the synthesis of proteins.

Even though the poor initiation context at the uORFs led to frequent leaky scanning in the 5′ leaders of DAP5 targets, our mutational analysis of start and STOP codons showed that translation of short uORFs was mandatory for expression of the main CDS. Consistent with a role in re-initiation, DAP5 function was also sensitive to the inhibition of termination and ribosome recycling. In addition, a subset of the DAP5 targets was less translated in cells deficient for DENR, an initiation/recycling factor previously implicated in the re-initiation of translation in animal cells (Ahmed et al., 2018; Bohlen et al., 2020; Castelo-Szekely et al., 2019; Schleich et al., 2014; Vasudevan et al., 2020).

Altogether, our work reports a previously unrecognized role for DAP5 in the control of translation in human cells.

### DAP5 regulates the synthesis of signalling proteins

Synthesis of developmental, regulatory and disease-relevant proteins often occurs on mRNAs with GC- and uORF-rich 5′ leaders (Fujii et al., 2017; Kozak, 1991; Renz et al., 2020; Wethmar et al., 2016). These intricate 5′ leaders are thought to limit the production of proteins that are detrimental to cells if overproduced or deregulated by imposing stringent translational control. Although the regulatory potential of these 5′ leaders has long been recognized, the molecular mechanisms enforcing translational control are largely unknown. We find that DAP5-dependent re-initiation is required for translation of the main CDS of mRNAs with 5′ leaders where structured regions and uORFs are abundant. DAP5 targets are enriched for mRNAs encoding components of different signalling pathways, such as kinases, phosphatases and GTPases, that control cell migration and adhesion, proliferation, differentiation and transcription. Among the DAP5 targets are members of WNT pathway (e.g. *WNT5A*, *GSK3B*, *TLE1/3/4*, *AXIN1*, *FZD8*, *EP300*, *LGR4*) with long and structured 5′ leaders (Nguyen et al., 2018), vascular endothelial growth factor signalling and MAPK cascade (e.g. *MAPK13*, *MAP2K2/3*, *MAP3K2*, *MAP4K4*), or different disease-associated genes and proto-oncogenes (e.g. *AKT1*, *CEBPA*, *RAF1*, *SKI*, *FYN*, *PIK3R2*). Thus, DAP5-dependent translational control of specific signalling components and enzymes that usually have dose-dependent functions efficiently regulates the overall strength of particular signalling pathways in response to stimuli, enabling cells to adapt or adopt different states according to the surrounding environment. Underscoring the physiological importance of DAP5 and re-initiation are the observations that DAP5 deletion in animals results in early embryonic lethality by blocking stem-cell differentiation (Nousch et al., 2007; Sugiyama et al., 2017; Takahashi et al., 2020; Yamanaka et al., 2000; Yoffe et al., 2016; Yoshikane et al., 2007). As several DAP5 targets are known oncogenes and disease-associated genes, future investigations are required to unveil the biological and functional implications of re-initiation in pathological settings. Together with the growing evidence that defective uORF function, polymorphisms and translational reprogramming at 5′ leaders contribute to various human diseases (Barbosa et al., 2013; Sendoel et al., 2017; Wethmar et al., 2016), our work opens new directions into whether uORF translation, re-initiation and DAP5 can be exploited for future therapeutic interventions.

Our results also highlight the functional importance of 5′ leaders, uORFs and re-initiation in the regulation of gene expression. A mechanistic understanding of the influence of alternative 5′ leaders, structured elements and the increased coding capacity of the genome as a consequence of re-initiation will provide exciting findings on how cells precisely tune protein expression levels.

## Acknowledgments

We dedicate this work to the memory of Elisa Izaurralde. We gratefully acknowledge that the study was conceived and carried out in her laboratory. We are thankful to Markus Landthaler and Ulrike Zinnall for their help with ribosome profiling, and Heike Budde for cloning of the pcDNA3.1-MCS-mCherry plasmid. This work was supported by the Max Planck Society.

## Author Contributions

R.W. designed and conducted the experiments assisted by M.-Y.C. L.K. performed luciferase assays and generated several constructs used in this study. I.H. assisted and contributed to the analysis of FACS data. E.V. contributed to data analysis. R.W. and C.I. conceived the project, interpreted the results, and wrote the manuscript. All authors read and corrected the manuscript.

The authors declare no competing interests.

## Materials and Methods

### Resource availability

Further information and requests for resources and reagents should be directed to and will be fulfilled by Catia Igreja.

### Materials availability

All unique/stable reagents generated in this study are available with a completed Materials Transfer Agreement.

### Experimental model and subject details

#### Cell lines

All cell lines were cultured at 37°C and 5% CO_2_ in Dulbecco’s Modified Eagle’s Medium (DMEM) supplemented with 10% fetal bovine serum, 2 mM Glutamine, 1x Penicillin and 1x Streptomycin.

### Methods details

#### DNA constructs

DNA constructs used in this study are listed in the Key Resources Table. All the constructs were confirmed by sequencing. The pCIneo-R-LUC, pEGFP-N3-F-LUC, pT7-EGFP-C1-MBP and pT7-EGFP-C1-4EBP plasmids were described previously (Lazzaretti et al., 2009; Peter et al., 2015b; Peter et al., 2019; Pillai et al., 2004). To produce the pT7-V5-SBP-C1-MBP, MBP cDNA was introduced in the XhoI and BamHI cut sites of the pT7-V5-SBP-C1 vector. *Hs* DAP5 and eIF4G cDNAs were introduced in the XhoI and KpnI or XhoI and HindIII restriction sites of the pT7-V5-SBP-C1 vector, respectively. To generate the *WNK1-*, *ROCK1-* and *AKT1-*R-LUC reporters, the respective 5′ leader sequences were obtained as synthetic cDNA clones from Invitrogen using the GeneArt tool. Using site-directed mutagenesis, an EcoRI restriction site was inserted upstream of the luciferase ORF present in the pCIneo-R-LUC vector, and EcoRI sites were removed from the *WNK1* and *ROCK1* 5′ leader sequences. The synthetic DNA strings were then inserted into the NheI and EcoRI restriction sites of the modified pCIneo-R-LUC vector. The pSFFV_mNG2(11)1-10 plasmid was a gift from Bo Huang (Addgene plasmid # 82610) (Feng et al., 2017) and the EBFP-N1 was a gift from Michael Davidson (Addgene plasmid # 54595). The mNG2 1-10 sequence present in pSFFV_mNG2(11)1-10 vector was subcloned into pDNA3.1 using the HindIII and BamHI restriction sites. To generate the EBFP fluorescent reporter with the *WNK1* 5′ leader, the EBFP cDNA was cloned between the EcoRI and XbaI restriction sites of a modified pCIneo-R-LUC, which contains an EcoRI cut site. In this cloning step, EBFP replaces R-LUC ORF. The *WNK1* 5′ leader was then inserted into the NheI and EcoRI sites of the pCIneo-EBFP vector. *WNK1* uORF2 was then replaced by the DNA sequence encoding mNG2 11^th^ β-strand using site-directed mutagenesis. The mCherry sequence was inserted into the XhoI and ApaI restriction sites of the pcDNA3.1-MCS vector. Human eRF1 coding sequence was cloned into the XhoI and HindIII sites of the pλN-HA-C1 vector (Clontech). The pλN-HA-C1-eRF1 plasmid was then used to generate the pλN-HA-C1-eRF1^AAQ^ (G183A G184A) dominant negative mutant by mutagenesis. Human eIF2β cDNA was cloned into the XhoI and BamHI sites of the pT7-EGFP-C1 vector. To generate the DAP5-eIF4G chimeras, DAP5 sequences corresponding to the MIF4G, MA3 and W2 domains were replaced by the respective eIF4G sequences. MIF4G: residues 738-998 of eIF4G iso9 replace residues 78-308 of DAP5. MA3: residues 1240-1435 of eIF4G iso9 replace residues 540-723 of DAP5. W2: residues 1444-1606 of eIF4G iso9 replace residues 730-907 of DAP5. To generate the reporters used in Figure 4, 5 and S5, the *WNK1* 5′ leader sequence was modified with the following nucleotide mutations. **uORF 118**: A391T; **uORF 30**: A391T, CCC467-469TGA; **uORF 19**: A391T, CGC434-436TGA; **uORF 9**: A391T, TCG404-406TAG; Δ**STOP IF**: G161C, A659T, A782T; Δ**STOP all F**: G31C, A96T, G161C, A213T, A391T, A444T, G570C, A659T, A733T, A782T, A864T, A868T; **uORF2+**: GTG93-95ATG, TGG111-113ATG; Δ**STOP1**: GTG93-95ATG, TGG111-113ATG, G161C; Δ**STOP1+2**: GTG93-95ATG, TGG111-113ATG, G161C, A659T; **NO STOP**: GTG93-95ATG, TGG111-113ATG, G161C, A659T, A782T; **uORF 49**: GTG93-95ATG, TGG111-113ATG, G161C, TCC240-242TGA; **uORF 39**: GTG93-95ATG, TGG111-113ATG, G161C, GTG210-212TAG, **uORF 29**: GTG93-95ATG, TGG111-113ATG, G161C, TCA180-182TGA. The plasmids expressing short hairpin RNAs (shRNAs) used in the knockdown experiments were derived from the pSUPERpuro plasmid (a gift from O. Mühlemann) containing the puromycin resistance gene for cell selection. The shRNA target sequences are listed in Table S4.

All the mutants used in this study were generated by site-directed mutagenesis using the QuickChange Site-Directed Mutagenesis kit (Stratagene).

#### Generation of the DAP5-null cell line

Two sgRNAs targeting DAP5 were designed and cloned into the pSpCas9(BB)-2A-Puro (PX459) vector [a gift from F. Zhang, Addgene plasmid 48139; (Ran et al., 2013)] using the CHOPCHOP (http://chopchop.cbu.uib.no) online tool as previously described (Peter et al., 2017). Briefly, HEK293T cells were transfected with the sgRNA-Cas9 vector. Forty-eight hours later, edited cells were selected with puromycin (3 μg/ml: Serva Electrophoresis). Serial dilutions in 96-well plates were used to isolate single cell clones. Genomic DNA was extracted from the different clones using the Wizard SV Genomic DNA Purification System (Promega). The DAP5 locus was PCR amplified and Sanger sequencing of the targeted genomic regions indicated two frameshift mutations in exon 10 (172 bp deletion in exon/intron 10, and a 1 bp insertion) targeted by sgDAP5-a. These mutations caused defective splicing and intron retention, as evidenced by subsequent RNA sequencing (Figure S1B). Two mutations were detected in exon 12 (1 bp insertion and 12 bp deletion) targeted by sgDAP5-b. The lack of DAP5 protein was further confirmed by western blotting (Figures 1D, S1C). RNA sequencing revealed that DAP5 transcript levels were severely reduced in the null cells compared to wild-type cells (Figure S1A), most likely as a result of non-sense mediated decay. The following guide sequences were used: sgDAP5-a: 5’-CACGTACCTTGGCTCGTTCA-3’; sgDAP5-b: 5’-ACACCATTGGGTTCCTCGCA-3’.

#### Ribosome profiling and RNA sequencing

For ribosome profiling and RNA sequencing HEK293T wild type and DAP5-null cells were plated on 10 cm dishes 24 hours before harvesting (3.2 x 10^6^ WT cells and 3.5 x 10^6^ null cells per plate). Cells were harvested as described in (Calviello et al., 2016). Importantly, cells were not incubated with cycloheximide before harvesting. Cycloheximide (100 μg/ml, Serva Electrophoresis) was only present in the washing and lysis buffer, as described in (Calviello et al., 2016). For total RNA sequencing, RNA was extracted using the RNeasy Mini Kit (50) (Qiagen) and processed according to the Illumina TruSeq RNA Sample Prep Kit. For ribosome profiling the original protocol (Ingolia et al., 2012) was used in a modified version also described in (Calviello et al., 2016). The ribosome profiling and total RNA sequencing pools were sequenced on an Illumina Hiseq3000 instrument. Reads originating from ribosomal RNA were removed using Bowtie2 (Langmead and Salzberg, 2012). Remaining reads of the RNA sequencing library were mapped onto the human genome using Tophat2 (Kim et al., 2013) which resulted in 15.7-20.5 million mapped reads with an overall read mapping rate >94% for the RNA sequencing experiment. Ribosome profiling reads were subjected to statistical analysis using RiboTaper that aims at identifying actively translating ribosomes based on the characteristic three-nucleotide periodicity (Calviello et al., 2016). Reads of 29 and 30 nucleotides length showed the best three-nucleotide periodicity and where therefore used for subsequent mapping onto the human genome. This resulted in 2.8-3.8 million mapped reads with an overall read mapping rate >95% for the ribosome profiling experiment. Read count analysis was performed using QuasR (Gaidatzis et al., 2015). Differential expression analysis was conducted using edgeR (McCarthy et al., 2012; Robinson et al., 2010). Translation efficiency (TE) was calculated using RiboDiff (Zhong et al., 2017).

Harringtonine and LTM datasets from human HEK293 cells were downloaded from the Sequence Read Archive database (accession: SRA056377). RocA and DENR datasets were retrieved from the GEO database accession numbers GSE70211 and GSE134020, respectively. Ribosomal RNA reads were filtered using Bowtie 2 (Langmead and Salzberg, 2012). The remaining reads were mapped on the hg19 (UCSC) human genome or the mm9 (UCSC) mouse genome with TopHat2 (Kim et al., 2013). No specific filters for read length were applied.

#### Analysis of GO terms and nucleotide compositions

Upregulated and downregulated gene groups were defined as being significantly deregulated (FDR<0.005) with a logFC>0 and logFC<0, respectively. No cut-off of the logFC value was applied so that genes with little but significant changes could also be detected. GO analysis was performed with the R based package goseq (Young et al., 2010). For analysis of 5′ leader nucleotide composition, the respective mRNA sequences were fetched using biomaRt (Durinck et al., 2005; Durinck et al., 2009). Analysis of GC content and length of 5′ leader was performed with R based scripts.

RNA structures were calculated using the ViennaRNA package 2.0 (Lorenz et al., 2011). Metagene analysis was performed using the Deeptools suite of functions (Ramirez et al., 2016). For uORF number, size and start codon analysis the accumulation of ribosome footprint on start codons was assessed using the ribosome profiling dataset in HEK293 cells treated with harringtonine (Lee et al., 2012). Identity of the start codon and the corresponding STOP codon was manually assigned.

Ribosome footprint density plots for individual sequencing tracks were visualized using the Integrative Genomics Viewer (IGV) visualization tool (Robinson et al., 2011; Thorvaldsdottir et al., 2013).

#### Transfections, northern and western blotting

In the rescue assays described in Figures 2, 3B, 4, 5 and 6, 0.64 x 10^6^ WT cells or 0.7 x 10^6^ null cells were transfected, after seeding in 6-well plates, using Lipofectamine 2000 (Invitrogen). The transfection mixtures contained different amounts of the plasmids expressing R-LUC, GFP-F-LUC or V5-SBP-fusion proteins (*WNK1*-R-LUC reporters: 0.5 μg; GFP-F-LUC: 0.25 μg ; V5-SBP-MBP: 0.3 μg; V5-SBP-DAP5 FL and MIF4G: 0.8 μg; V5-SBP-DAP5 eIF4A*: 3.25 μg; V5-SBP-DAP5 ΔW2: 1.2 μg; V5-SBP-eIF4G FL: 3.25 μg; V5-SBP-eIF4G ΔN: 0.8 μg; V5-SBP-Chimeras: 0.8 μg). For the experiment shown in Figure 7, λN-HA-eRF1 G183A G184A was titrated using 0.25 μg; 0.75 μg; and 1.25 μg of plasmid DNA.

Cells were harvested two days after transfection and firefly and *Renilla* luciferase activities were measured using the Dual-Luciferase reporter assay system (Promega). Total RNA was isolated using TriFast (Peqlab biotechnologies). For northern blotting, total RNA was separated in 2% glyoxal agarose gels and blotted onto a positively charged nylon membrane (GeneScreen Plus, Perkin Elmer). [^32^P]-labelled probes specific for each transcript were generated by linear PCR. Hybridizations were carried out in hybridization solution (0.5 M NaP pH=7.0, 7% SDS, 1 mM EDTA pH=8.0) at 65°C overnight. After extensive washes with washing solution (40 mM NaP pH=7.0, 1% SDS, 1 mM EDTA pH=8.0), the membranes were exposed and band intensities were quantified by PhosphoImager.

Western blot was performed using standard methods. In brief, cells were washed with PBS and lysed with sample buffer (100 mM Tris-HCl pH=6.8, 4% SDS, 20% glycerol, 0.2 M DTT) followed by boiling 5 minutes at 95°C and vortexing to shear genomic DNA. After SDS-PAGE, proteins were transferred onto a nitrocellulose membrane (Santa Cruz Biotechnology) by tank transfer. Primary antibodies were incubated overnight at 4°C and secondary antibodies for an hour at room temperature. All western blots were developed with freshly mixed 10A: 1B ECL solutions and 0.01% H_2_O_2_ [Solution A: 0.025 % Luminol (Roth) in 0.1 M Tris-HCl pH=8.6; Solution B: 0.11% P-Coumaric acid (Sigma Aldrich) in DMSO]. Antibodies used in this study are listed in the Key Resource Table.

#### Reverse transcription (RT) and quantitative PCR (qPCR)

1 µg of RNA was mixed with 0.66 μg of random hexamer primers (N_6_) and denatured at 72°C for 5 min. After addition of a reaction mixture containing a final concentration of 1 x RT buffer, 20 U RiboLock RNase Inhibitor (Thermo Scientific) and 1 mM dNTPs, the RNA samples were incubated at 37°C for 5 min. Incubation with RevertAid H Minus Reverse Transcriptase (200 U, Thermo Scientific) was first performed for 10 min at 25°C, and then at 42°C for one hour. The RT reaction was stopped by incubating the samples for 10 min at 70°C. The qPCR was performed with 1 x iTaq SYBR Green Supermix (Biorad), 0.4 µM of each primer and 1 µl of the cDNA sample. mRNA levels were determined by qPCR using sequence-specific primers for the indicated transcripts. qPCR primers designed using Primer-BLAST (NCBI) are listed in Table S4. Normalized transcript expression ratios from three independent experiments were determined using the Livak method (Livak and Schmittgen, 2001).

#### Polysome profiling

Polysome profiles were performed as described before (Kuzuoglu-Ozturk et al., 2016). HEK293T cells were pre-treated with cycloheximide (50 µg/ml) for 30 min. Lysates were prepared in lysis buffer (10 mM Tris-HCl pH=7.4, 10 mM NaCl, 1.5 mM MgCl_2_, 0.5% Triton X-100, 2 mM DTT, 50 µg/ml cycloheximide) and polysomes separated on a 10-50% sucrose gradient in gradient buffer (10 mM Tris-HCl pH 7.4, 75 mM KCl, 1.5 mM MgCl_2_). Polysome fractions were collected using the Teledyne Isco Density Gradient Fractionation System.

To isolate RNA from sucrose fractions, samples were first digested with proteinase K (Sigma Aldrich, 1% of the sample volume; 100 mg/ml in 50 mM Tris-HCl pH=8.1, 10 mM CaCl_2_ buffer) at 37°C for 45 min and shaking at 400 rpm. The digested samples were mixed with 1 volume of Phenol:Chloroform:Isoamyl alcohol (PanReac AppliChem, 25:24:1, v/v), vortexed and spun down 5 min at 20,000 *g* at 4°C. Supernatants were transferred into 3 volumes of 100% ethanol, 0.1 volumes of 3M NaOAc pH=5.2 and 1 µl of GlycoBlue, and precipitated at -20°C. Samples were pelleted for 30 min at 20,000 *g* and 4°C, washed once with 100% ethanol and another time with 70% ethanol, dried and resuspended in 30 µl H_2_O. Fractions were reverse transcribed and analysed by RT-qPCR.

#### RNA pulldown

For the RNA pulldown, 3 x 10^6^ WT HEK293T cells were plated in 10 cm plates and transfected using Lipofectamine 2000 (Invitrogen) with the following plasmids expressing V5-SBP fusions: MBP (1.5 μg), DAP5 FL (4 μg) and eIF4A* (15 μg), MIF4G (4 μg) or ΔW2 (6 μg mutants, eIF4G FL (15 μg) or ΔN (4 μg), and Chimeras (4 μg). A detailed description of the RNA immunoprecipitation procedure can be found in (Kuzuoglu-Ozturk et al., 2016). Cells were harvested 48 hours post transfection, washed with ice cold PBS and lysed on ice for 15 minutes in 500 μl of NET buffer [50 mM Tris-HCl pH=7.5, 150 mM NaCl, 0.1% Triton X-100, 1 mM EDTA pH=8.0, 10 % glycerol, supplemented with 1x protease inhibitors (Roche)]. Cell debris was removed by centrifugation at 16,000 *g* and 4°C. Input samples (5% of the total) were collected for western blotting and RT-qPCR. Cell lysates were immediately incubated with 50 μl of a 50% slurry of streptavidin beads pre-incubated with yeast RNA (250 μg of yeast RNA/100 μl of 50% slurry). Beads were washed 3 times with NET buffer and resuspended in 1 ml of NET buffer without detergent. An aliquot (20% of the total) of the bead suspension was mixed with SDS-PAGE sample buffer for western blotting after centrifugation to pellet the resin. The remaining beads were used for RNA isolation with TriFast (Peqlab Biotechnologies). cDNA of the input and precipitated fractions (20% each) was prepared and analysed using qPCR (5% of the cDNA), as described above. The list of primers used for the qPCR experiments can be found in Table S4.

#### Pulldown assays

Pulldown assays were performed in the presence of RNase A as described previously (Peter et al., 2015). HEK293T cells were grown in 10 cm dishes and transfected using Lipofectamine 2000 (Invitrogen) according to the manufacturer’s recommendations. The transfection mixtures in Figures S4A and B contained 1.5 μg of V5-SBP-MBP, 4 μg of V5-SBP-DAP5, and 5 μg of GFP-eIF2β. After transfection, cells were treated as described in the RNA pulldown section, with the exception that the streptavidin beads were not incubated with yeast RNA and the samples were solely used for immunoblotting.

For the cap pulldown transfection mixtures contained 1 μg GFP-MBP or 12 μ GFP-chimeric-4EBP. Cap-bound proteins were pulled down using γ-Aminophenyl-m^7^GTP beads (Jena Bioscience).

#### Flow cytometry

Cells were seeded (0.6 x 10^6^ WT and 0.7 x 10^6^ DAP5-null HEK293T cells) in 6-well plates 24 hours before transfection. Transfections were carried out with Lipofectamine 2000 (Invitrogen), with the following transfection mixtures: *WNK1*-mNG2_11_-EBFP (0.35 µg), mNG2_1-10_ (1 µg), mCherry (10 ng), V5-SBP-MBP (0.25 µg) or DAP5 (0.65 µg). 48 hours after transfection, cells were trypsinized, sedimented (1000 rpm for 3 min at room temperature), resuspended in 1% FBS in PBS, and analysed using the Becton Dickinson FACSMelody^™^ Cell Sorter and FlowJo software (Becton Dickison). To determine mNG2, EBFP and mCherry positive events, we analysed non-transfected and control transfected cells. Cut-offs were applied uniformly for all measured conditions.

#### Knockdowns

0.64 x 10^6^ HEK293T cells were transfected with 2 µg pSUPER-puro scramble control or ABCE1, MCTS1, DENR and eIF2D shRNAs, after seeding in 6-well plates, using Lipofectamine 2000 (Invitrogen). 24 hours after transfection cells were treated with 5 µM puromycin (Serva Electrophoresis) for 24 hours. Selected cells were re-seeded and re-transfected with DNA mixtures containing 0.5 µg of *WNK1*-R-LUC reporter plasmid and 0.25 µg of the control GFP-F-LUC reporter.

### Quantification and statistical analyses

Figures 1B, S1D. Upregulated and downregulated genes were identified using log_2_Fold Change (FC) between null and control cells > 0 or < 0, respectively, and False Discovery Rates (FDR) < 0.005.

Figure 1C. The quantitative value represented in the graphs corresponds to -log_10_(q-value) determined by the GOseq analysis tool (Young et al., 2010).

Figures 1H, I. Boxes indicate the 25^th^ to 75^th^ percentiles; black line inside the box represents the median; whiskers indicate the extend of the highest and lowest observations; dots show the outliers.

Figures 6J, 6K. Boxes represent the 25^th^ to 75^th^ percentiles; black line shows the median and the cross the average; whiskers show the variability outside the upper and lower quartiles; dots show the outliers. Significance determined by one-sided Wilcoxon rank-sum test and indicated if p<2.2e^-16^ (**).

Figure 1J. The quantitative values represented in the pie chart indicate the percentage of uORFs containing canonical and near-cognate start codons or other codon sequence in the 5′ leaders of the DAP5 targets.

Figures 2, 3, 4, 5, 7, S1J-L, S3E-G, S4F, S4G, S5, S6, S7. The quantitative value that is graphed represents the mean mRNA or protein level values; error bars represent standard deviations from three independent experiments. In the RT-qPCR experiments, normalized transcript expression ratios from three independent experiments were determined using the Livak method (Livak and Schmittgen, 2001).

Figures S3A-D. Length, GC content, minimum free energy and TE were determined for the 5′ leaders of DAP5 targets and in all other mRNAs expressed in HEK293T cells. Statistical significance was calculated with the Wilcoxon rank sum test.

Figures S4C, S7H. The hypergeometric test (phyper) in R was applied to estimate the likelihood of list overlap.

## Notes

### Competing Interest Statement

The authors have declared no competing interest.

## References

Ahmed, Y.L., Schleich, S., Bohlen, J., Mandel, N., Simon, B., Sinning, I., and Teleman, A.A. (2018). DENR-MCTS1 heterodimerization and tRNA recruitment are required for translation reinitiation. PLoS Biol 16, e2005160.

Barbosa, C., Peixeiro, I., and Romao, L. (2013). Gene expression regulation by upstream open reading frames and human disease. PLoS Genet 9, e1003529.

Bazzini, A.A., Johnstone, T.G., Christiano, R., Mackowiak, S.D., Obermayer, B., Fleming, E.S., Vejnar, C.E., Lee, M.T., Rajewsky, N., Walther, T.C., et al. (2014). Identification of small ORFs in vertebrates using ribosome footprinting and evolutionary conservation. EMBO J 33, 981–993.

Bohlen, J., Harbrecht, L., Blanco, S., Clemm von Hohenberg, K., Fenzl, K., Kramer, G., Bukau, B., and Teleman, A.A. (2020). DENR promotes translation reinitiation via ribosome recycling to drive expression of oncogenes including ATF4. Nat Commun 11, 4676.

Brown, A., Shao, S., Murray, J., Hegde, R.S., and Ramakrishnan, V. (2015). Structural basis for stop codon recognition in eukaryotes. Nature 524, 493–496.

Bukhari, S.I.A., Truesdell, S.S., Lee, S., Kollu, S., Classon, A., Boukhali, M., Jain, E., Mortensen, R.D., Yanagiya, A., Sadreyev, R.I., et al. (2016). A Specialized Mechanism of Translation Mediated by FXR1a-Associated MicroRNP in Cellular Quiescence. Mol Cell 61, 760–773.

Calviello, L., Mukherjee, N., Wyler, E., Zauber, H., Hirsekorn, A., Selbach, M., Landthaler, M., Obermayer, B., and Ohler, U. (2016). Detecting actively translated open reading frames in ribosome profiling data. Nat Methods 13, 165–170.

Castelo-Szekely, V., De Matos, M., Tusup, M., Pascolo, S., Ule, J., and Gatfield, D. (2019). Charting DENR-dependent translation reinitiation uncovers predictive uORF features and links to circadian timekeeping via Clock. Nucleic Acids Res 47, 5193–5209.

Chen, J., Brunner, A.D., Cogan, J.Z., Nunez, J.K., Fields, A.P., Adamson, B., Itzhak, D.N., Li, J.Y., Mann, M., Leonetti, M.D., et al. (2020). Pervasive functional translation of noncanonical human open reading frames. Science 367, 1140–1146.

de la Parra, C., Ernlund, A., Alard, A., Ruggles, K., Ueberheide, B., and Schneider, R.J. (2018). A widespread alternate form of cap-dependent mRNA translation initiation. Nat Commun 9, 3068.

Durinck, S., Moreau, Y., Kasprzyk, A., Davis, S., De Moor, B., Brazma, A., and Huber, W. (2005). BioMart and Bioconductor: a powerful link between biological databases and microarray data analysis. Bioinformatics 21, 3439–3440.

Durinck, S., Spellman, P.T., Birney, E., and Huber, W. (2009). Mapping identifiers for the integration of genomic datasets with the R/Bioconductor package biomaRt. Nat Protoc 4, 1184–1191.

Feng, S., Sekine, S., Pessino, V., Li, H., Leonetti, M.D., and Huang, B. (2017). Improved split fluorescent proteins for endogenous protein labeling. Nat Commun 8, 370.

Fritsch, C., Herrmann, A., Nothnagel, M., Szafranski, K., Huse, K., Schumann, F., Schreiber, S., Platzer, M., Krawczak, M., Hampe, J., et al. (2012). Genome-wide search for novel human uORFs and N-terminal protein extensions using ribosomal footprinting. Genome Res 22, 2208–2218.

Frolova, L.Y., Tsivkovskii, R.Y., Sivolobova, G.F., Oparina, N.Y., Serpinsky, O.I., Blinov, V.M., Tatkov, S.I., and Kisselev, L.L. (1999). Mutations in the highly conserved GGQ motif of class 1 polypeptide release factors abolish ability of human eRF1 to trigger peptidyl-tRNA hydrolysis. RNA 5, 1014–1020.

Fujii, K., Shi, Z., Zhulyn, O., Denans, N., and Barna, M. (2017). Pervasive translational regulation of the cell signalling circuitry underlies mammalian development. Nat Commun 8, 14443.

Gaidatzis, D., Lerch, A., Hahne, F., and Stadler, M.B. (2015). QuasR: quantification and annotation of short reads in R. Bioinformatics 31, 1130–1132.

Haizel, S.A., Bhardwaj, U., Gonzalez, R.L., Jr., Mitra, S., and Goss, D.J. (2020). 5’-UTR recruitment of the translation initiation factor eIF4GI or DAP5 drives cap-independent translation of a subset of human mRNAs. J Biol Chem 295, 11693–11706.

Hashem, Y., and Frank, J. (2018). The Jigsaw Puzzle of mRNA Translation Initiation in Eukaryotes: A Decade of Structures Unraveling the Mechanics of the Process. Annu Rev Biophys 47, 125–151.

Henis-Korenblit, S., Shani, G., Sines, T., Marash, L., Shohat, G., and Kimchi, A. (2002). The caspase-cleaved DAP5 protein supports internal ribosome entry site-mediated translation of death proteins. Proc Natl Acad Sci U S A 99, 5400–5405.

Henis-Korenblit, S., Strumpf, N.L., Goldstaub, D., and Kimchi, A. (2000). A novel form of DAP5 protein accumulates in apoptotic cells as a result of caspase cleavage and internal ribosome entry site-mediated translation. Mol Cell Biol 20, 496–506.

Hundsdoerfer, P., Thoma, C., and Hentze, M.W. (2005). Eukaryotic translation initiation factor 4GI and p97 promote cellular internal ribosome entry sequence-driven translation. Proc Natl Acad Sci U S A 102, 13421–13426.

Imataka, H., Gradi, A., and Sonenberg, N. (1998). A newly identified N-terminal amino acid sequence of human eIF4G binds poly(A)-binding protein and functions in poly(A)-dependent translation. EMBO J 17, 7480–7489.

Imataka, H., Olsen, H.S., and Sonenberg, N. (1997). A new translational regulator with homology to eukaryotic translation initiation factor 4G. EMBO J 16, 817–825.

Ingolia, N.T. (2014). Ribosome profiling: new views of translation, from single codons to genome scale. Nat Rev Genet 15, 205–213.

Ingolia, N.T., Brar, G.A., Rouskin, S., McGeachy, A.M., and Weissman, J.S. (2012). The ribosome profiling strategy for monitoring translation in vivo by deep sequencing of ribosome-protected mRNA fragments. Nat Protoc 7, 1534–1550.

Ingolia, N.T., Lareau, L.F., and Weissman, J.S. (2011). Ribosome profiling of mouse embryonic stem cells reveals the complexity and dynamics of mammalian proteomes. Cell 147, 789–802.

Iwasaki, S., Floor, S.N., and Ingolia, N.T. (2016). Rocaglates convert DEAD-box protein eIF4A into a sequence-selective translational repressor. Nature 534, 558–561.

Jackson, R.J., Hellen, C.U., and Pestova, T.V. (2012). Termination and post-termination events in eukaryotic translation. Adv Protein Chem Struct Biol 86, 45–93.

Jonas, S., Weichenrieder, O., and Izaurralde, E. (2013). An unusual arrangement of two 14-3-3-like domains in the SMG5-SMG7 heterodimer is required for efficient nonsense-mediated mRNA decay. Genes Dev 27, 211–225.

Kara, G., Tuncer, S., Turk, M., and Denkbas, E.B. (2015). Downregulation of ABCE1 via siRNA affects the sensitivity of A549 cells against chemotherapeutic agents. Med Oncol 32, 103.

Kim, D., Pertea, G., Trapnell, C., Pimentel, H., Kelley, R., and Salzberg, S.L. (2013). TopHat2: accurate alignment of transcriptomes in the presence of insertions, deletions and gene fusions. Genome Biol 14, R36.

Kozak, M. (1990). Downstream secondary structure facilitates recognition of initiator codons by eukaryotic ribosomes. Proc Natl Acad Sci U S A 87, 8301–8305.

Kozak, M. (1991). An analysis of vertebrate mRNA sequences: intimations of translational control. J Cell Biol 115, 887–903.

Kozak, M. (2002). Pushing the limits of the scanning mechanism for initiation of translation. Gene 299, 1–34.

Kuzuoglu-Ozturk, D., Bhandari, D., Huntzinger, E., Fauser, M., Helms, S., and Izaurralde, E. (2016). miRISC and the CCR4-NOT complex silence mRNA targets independently of 43S ribosomal scanning. EMBO J 35, 1186–1203.

Lamphear, B.J., Kirchweger, R., Skern, T., and Rhoads, R.E. (1995). Mapping of functional domains in eukaryotic protein synthesis initiation factor 4G (eIF4G) with picornaviral proteases. Implications for cap-dependent and cap-independent translational initiation. J Biol Chem 270, 21975–21983.

Langmead, B., and Salzberg, S.L. (2012). Fast gapped-read alignment with Bowtie 2. Nat Methods 9, 357–359.

Lazzaretti, D., Tournier, I., and Izaurralde, E. (2009). The C-terminal domains of human TNRC6A, TNRC6B, and TNRC6C silence bound transcripts independently of Argonaute proteins. RNA 15, 1059–1066.

Lee, S., Liu, B., Lee, S., Huang, S.X., Shen, B., and Qian, S.B. (2012). Global mapping of translation initiation sites in mammalian cells at single-nucleotide resolution. Proc Natl Acad Sci U S A 109, E2424–2432.

Lee, S.H., and McCormick, F. (2006). p97/DAP5 is a ribosome-associated factor that facilitates protein synthesis and cell proliferation by modulating the synthesis of cell cycle proteins. EMBO J 25, 4008–4019.

Leonetti, M.D., Sekine, S., Kamiyama, D., Weissman, J.S., and Huang, B. (2016). A scalable strategy for high-throughput GFP tagging of endogenous human proteins. Proc Natl Acad Sci U S A 113, E3501–3508.

Lewis, S.M., Cerquozzi, S., Graber, T.E., Ungureanu, N.H., Andrews, M., and Holcik, M. (2008). The eIF4G homolog DAP5/p97 supports the translation of select mRNAs during endoplasmic reticulum stress. Nucleic Acids Res 36, 168–178.

Liberman, N., Gandin, V., Svitkin, Y.V., David, M., Virgili, G., Jaramillo, M., Holcik, M., Nagar, B., Kimchi, A., and Sonenberg, N. (2015). DAP5 associates with eIF2beta and eIF4AI to promote Internal Ribosome Entry Site driven translation. Nucleic Acids Res 43, 3764–3775.

Liberman, N., Marash, L., and Kimchi, A. (2009). The translation initiation factor DAP5 is a regulator of cell survival during mitosis. Cell cycle 8, 204–209.

Livak, K.J., and Schmittgen, T.D. (2001). Analysis of relative gene expression data using real-time quantitative PCR and the 2(-Delta Delta C(T)) Method. Methods 25, 402–408.

Lorenz, R., Bernhart, S.H., Honer Zu Siederdissen, C., Tafer, H., Flamm, C., Stadler, P.F., and Hofacker, I.L. (2011). ViennaRNA Package 2.0. Algorithms Mol Biol 6, 26.

Mader, S., Lee, H., Pause, A., and Sonenberg, N. (1995). The translation initiation factor eIF-4E binds to a common motif shared by the translation factor eIF-4 gamma and the translational repressors 4E-binding proteins. Mol Cell Biol 15, 4990–4997.

Marash, L., and Kimchi, A. (2005). DAP5 and IRES-mediated translation during programmed cell death. Cell Death Differ 12, 554–562.

Marash, L., Liberman, N., Henis-Korenblit, S., Sivan, G., Reem, E., Elroy-Stein, O., and Kimchi, A. (2008). DAP5 promotes cap-independent translation of Bcl-2 and CDK1 to facilitate cell survival during mitosis. Mol Cell 30, 447–459.

McCarthy, D.J., Chen, Y., and Smyth, G.K. (2012). Differential expression analysis of multifactor RNA-Seq experiments with respect to biological variation. Nucleic Acids Res 40, 4288–4297.

Merrick, W.C., and Pavitt, G.D. (2018). Protein Synthesis Initiation in Eukaryotic Cells. Cold Spring Harb Perspect Biol 10.

Nguyen, T.M., Kabotyanski, E.B., Dou, Y., Reineke, L.C., Zhang, P., Zhang, X.H., Malovannaya, A., Jung, S.Y., Mo, Q., Roarty, K.P., et al. (2018). FGFR1-Activated Translation of WNT Pathway Components with Structured 5’ UTRs Is Vulnerable to Inhibition of EIF4A-Dependent Translation Initiation. Cancer Res 78, 4229–4240.

Nousch, M., Reed, V., Bryson-Richardson, R.J., Currie, P.D., and Preiss, T. (2007). The eIF4G-homolog p97 can activate translation independent of caspase cleavage. RNA 13, 374–384.

Pelletier, J., and Sonenberg, N. (2019). The Organizing Principles of Eukaryotic Ribosome Recruitment. Annu Rev Biochem 88, 307–335.

Peter, D., Igreja, C., Weber, R., Wohlbold, L., Weiler, C., Ebertsch, L., Weichenrieder, O., and Izaurralde, E. (2015). Molecular architecture of 4E-BP translational inhibitors bound to eIF4E. Mol Cell 57, 1074–1087.

Peter, D., Ruscica, V., Bawankar, P., Weber, R., Helms, S., Valkov, E., Igreja, C., and Izaurralde, E. (2019). Molecular basis for GIGYF-Me31B complex assembly in 4EHP- mediated translational repression. Genes Dev 33, 1355–1360.

Pillai, R.S., Artus, C.G., and Filipowicz, W. (2004). Tethering of human Ago proteins to mRNA mimics the miRNA-mediated repression of protein synthesis. Rna 10, 1518–1525.

Prevot, D., Decimo, D., Herbreteau, C.H., Roux, F., Garin, J., Darlix, J.L., and Ohlmann, T. (2003). Characterization of a novel RNA-binding region of eIF4GI critical for ribosomal scanning. EMBO J 22, 1909–1921.

Pyronnet, S., Imataka, H., Gingras, A.C., Fukunaga, R., Hunter, T., and Sonenberg, N. (1999). Human eukaryotic translation initiation factor 4G (eIF4G) recruits mnk1 to phosphorylate eIF4E. EMBO J 18, 270–279.

Ramirez, F., Ryan, D.P., Gruning, B., Bhardwaj, V., Kilpert, F., Richter, A.S., Heyne, S., Dundar, F., and Manke, T. (2016). deepTools2: a next generation web server for deep-sequencing data analysis. Nucleic Acids Res 44, W160–165.

Ran, F.A., Hsu, P.D., Wright, J., Agarwala, V., Scott, D.A., and Zhang, F. (2013). Genome engineering using the CRISPR-Cas9 system. Nat Protoc 8, 2281–2308.

Renz, P.F., Valdivia-Francia, F., and Sendoel, A. (2020). Some like it translated: small ORFs in the 5’UTR. Exp Cell Res 396, 112229.

Robinson, J.T., Thorvaldsdottir, H., Winckler, W., Guttman, M., Lander, E.S., Getz, G., and Mesirov, J.P. (2011). Integrative genomics viewer. Nat Biotechnol 29, 24–26.

Robinson, M.D., McCarthy, D.J., and Smyth, G.K. (2010). edgeR: a Bioconductor package for differential expression analysis of digital gene expression data. Bioinformatics 26, 139–140.

Rodan, A.R., and Jenny, A. (2017). WNK Kinases in Development and Disease. Curr Top Dev Biol 123, 1–47.

Scheper, G.C., Morrice, N.A., Kleijn, M., and Proud, C.G. (2001). The mitogen-activated protein kinase signal-integrating kinase Mnk2 is a eukaryotic initiation factor 4E kinase with high levels of basal activity in mammalian cells. Mol Cell Biol 21, 743–754.

Schleich, S., Acevedo, J.M., Clemm von Hohenberg, K., and Teleman, A.A. (2017). Identification of transcripts with short stuORFs as targets for DENR*MCTS1-dependent translation in human cells. Sci Rep 7, 3722.

Schleich, S., Strassburger, K., Janiesch, P.C., Koledachkina, T., Miller, K.K., Haneke, K., Cheng, Y.S., Kuechler, K., Stoecklin, G., Duncan, K.E., et al. (2014). DENR-MCT-1 promotes translation re-initiation downstream of uORFs to control tissue growth. Nature 512, 208–212.

Sendoel, A., Dunn, J.G., Rodriguez, E.H., Naik, S., Gomez, N.C., Hurwitz, B., Levorse, J., Dill, B.D., Schramek, D., Molina, H., et al. (2017). Translation from unconventional 5’ start sites drives tumour initiation. Nature 541, 494–499.

Shao, S., Murray, J., Brown, A., Taunton, J., Ramakrishnan, V., and Hegde, R.S. (2016). Decoding Mammalian Ribosome-mRNA States by Translational GTPase Complexes. Cell 167, 1229–1240 e1215.

Skabkin, M.A., Skabkina, O.V., Dhote, V., Komar, A.A., Hellen, C.U., and Pestova, T.V. (2010). Activities of Ligatin and MCT-1/DENR in eukaryotic translation initiation and ribosomal recycling. Genes Dev 24, 1787–1801.

Sugiyama, H., Takahashi, K., Yamamoto, T., Iwasaki, M., Narita, M., Nakamura, M., Rand, T.A., Nakagawa, M., Watanabe, A., and Yamanaka, S. (2017). Nat1 promotes translation of specific proteins that induce differentiation of mouse embryonic stem cells. Proc Natl Acad Sci U S A 114, 340–345.

Takahashi, K., Jeong, D., Wang, S., Narita, M., Jin, X., Iwasaki, M., Perli, S.D., Conklin, B.R., and Yamanaka, S. (2020). Critical Roles of Translation Initiation and RNA Uridylation in Endogenous Retroviral Expression and Neural Differentiation in Pluripotent Stem Cells. Cell Rep 31, 107715.

Thorvaldsdottir, H., Robinson, J.T., and Mesirov, J.P. (2013). Integrative Genomics Viewer (IGV): high-performance genomics data visualization and exploration. Brief Bioinform 14, 178–192.

Topisirovic, I., Svitkin, Y.V., Sonenberg, N., and Shatkin, A.J. (2011). Cap and cap-binding proteins in the control of gene expression. Wiley Interdiscip Rev RNA 2, 277–298.

Vasudevan, D., Neuman, S.D., Yang, A., Lough, L., Brown, B., Bashirullah, A., Cardozo, T., and Ryoo, H.D. (2020). Translational induction of ATF4 during integrated stress response requires noncanonical initiation factors eIF2D and DENR. Nat Commun 11, 4677.

Virgili, G., Frank, F., Feoktistova, K., Sawicki, M., Sonenberg, N., Fraser, C.S., and Nagar, B. (2013). Structural analysis of the DAP5 MIF4G domain and its interaction with eIF4A. Structure 21, 517–527.

Weingarten-Gabbay, S., Khan, D., Liberman, N., Yoffe, Y., Bialik, S., Das, S., Oren, M., and Kimchi, A. (2014). The translation initiation factor DAP5 promotes IRES-driven translation of p53 mRNA. Oncogene 33, 611–618.

Wethmar, K., Schulz, J., Muro, E.M., Talyan, S., Andrade-Navarro, M.A., and Leutz, A. (2016). Comprehensive translational control of tyrosine kinase expression by upstream open reading frames. Oncogene 35, 1736–1742.

Yamanaka, S., Zhang, X.Y., Maeda, M., Miura, K., Wang, S., Farese, R.V., Jr., Iwao, H., and Innerarity, T.L. (2000). Essential role of NAT1/p97/DAP5 in embryonic differentiation and the retinoic acid pathway. EMBO J 19, 5533–5541.

Yanagiya, A., Svitkin, Y.V., Shibata, S., Mikami, S., Imataka, H., and Sonenberg, N. (2009). Requirement of RNA binding of mammalian eukaryotic translation initiation factor 4GI (eIF4GI) for efficient interaction of eIF4E with the mRNA cap. Mol Cell Biol 29, 1661–1669.

Yoffe, Y., David, M., Kalaora, R., Povodovski, L., Friedlander, G., Feldmesser, E., Ainbinder, E., Saada, A., Bialik, S., and Kimchi, A. (2016). Cap-independent translation by DAP5 controls cell fate decisions in human embryonic stem cells. Genes Dev 30, 1991–2004.

Yoshikane, N., Nakamura, N., Ueda, R., Ueno, N., Yamanaka, S., and Nakamura, M. (2007). Drosophila NAT1, a homolog of the vertebrate translational regulator NAT1/DAP5/p97, is required for embryonic germband extension and metamorphosis. Dev Growth Differ 49, 623–634.

Young, D.J., Makeeva, D.S., Zhang, F., Anisimova, A.S., Stolboushkina, E.A., Ghobakhlou, F., Shatsky, I.N., Dmitriev, S.E., Hinnebusch, A.G., and Guydosh, N.R. (2018). Tma64/eIF2D, Tma20/MCT-1, and Tma22/DENR Recycle Post-termination 40S Subunits In Vivo. Mol Cell 71, 761–774 e765.

Young, M.D., Wakefield, M.J., Smyth, G.K., and Oshlack, A. (2010). Gene ontology analysis for RNA-seq: accounting for selection bias. Genome Biol 11, R14.

Zhang, P., Chen, X.B., Ding, B.Q., Liu, H.L., and He, T. (2018). Down-regulation of ABCE1 inhibits temozolomide resistance in glioma through the PI3K/Akt/NF-kappaB signaling pathway. Biosci Rep 38.

Zhong, Y., Karaletsos, T., Drewe, P., Sreedharan, V.T., Kuo, D., Singh, K., Wendel, H.G., and Ratsch, G. (2017). RiboDiff: detecting changes of mRNA translation efficiency from ribosome footprints. Bioinformatics 33, 139–141.

