## Supplemental Figures and captions for "DAP5 enables translation re-initiation on structured messenger RNAs"

**Weber *et al.* Figure S1**

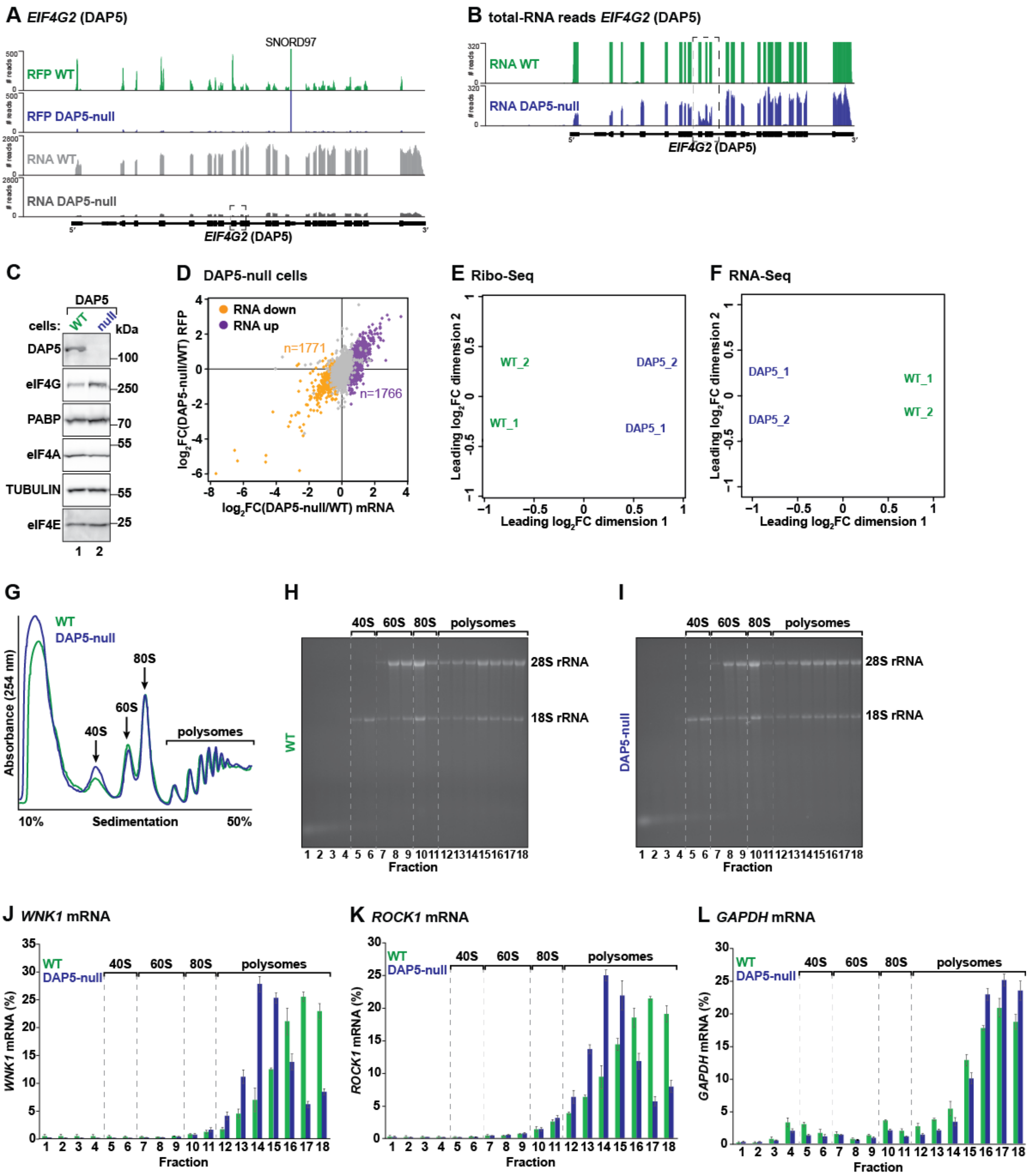


### Figure S1, related to Figure 1. Characterization of DAP5-null cells

**(A, B)** Ribosome footprints and total mRNA reads distribution along *WNK1* mRNA in wild type (WT) and DAP5-null cells. Of note, RFP and total RNA counts for *WNK1* are drastically reduced in the null cells. Dashed box indicates the position of the CRISPR-Cas9 edited region. In panel B, read counts scale for total RNA in DAP5-null cells was changed to show the presence of reads in intron 9 as a result of genome editing. SNORD97 (small nucleolar RNA, C/D Box 97) is encoded in intron 13 of *DAP5*.

**(C)** Western blot demonstrating the loss of DAP5 expression in the null cells. The expression of eIF4E, eIF4G and eIF4A is not decreased in the absence of DAP5. eIF4G and PABP protein levels are even increased in the null cells. TUBULIN served as a loading control. TUBULIN antibody recognizes an epitope common among the $\alpha$-tubulin subunits.

**(D)** Comparative analysis of translation efficiency (TE) in wild type (WT) and DAP5-null HEK293T cells as described in Figure 1A. Genes with increased (n=1771) and decreased (n=1766) mRNA abundance are highlighted in purple and orange, respectively.

**(E, F)** Multidimensional scaling (MDS) analysis for the Ribo-Seq (E) and RNA-Seq (F) replicate libraries from HEK293T wild type (WT) and DAP5-null cells.

**(G)** UV absorbance profile at 254 nm of HEK293T WT (green) and DAP5-null cells (blue) cell extracts after polysome sedimentation in a sucrose gradient. Absorbance peaks at 254 nm representing free 40S and 60S subunits, 80S monosomes and polysomes are indicated.

**(H, I)** Ethidium bromide staining of total RNA extracted from the different sucrose fractions. Due to high abundance, 28S and 18S rRNAs positions in the gel are readily detected.

**(J-L)** Abundance profiles for *WNK1* (J), *ROCK1* (K) and *GAPDH* (L) mRNAs across the density gradient in WT (green) and DAP5-null (blue) cells. mRNA abundance was determined by quantitative PCR (qPCR). Bars represent the mean value; error bars represent standard deviations (SD) (n=3).

Weber *et al.* Figure S2

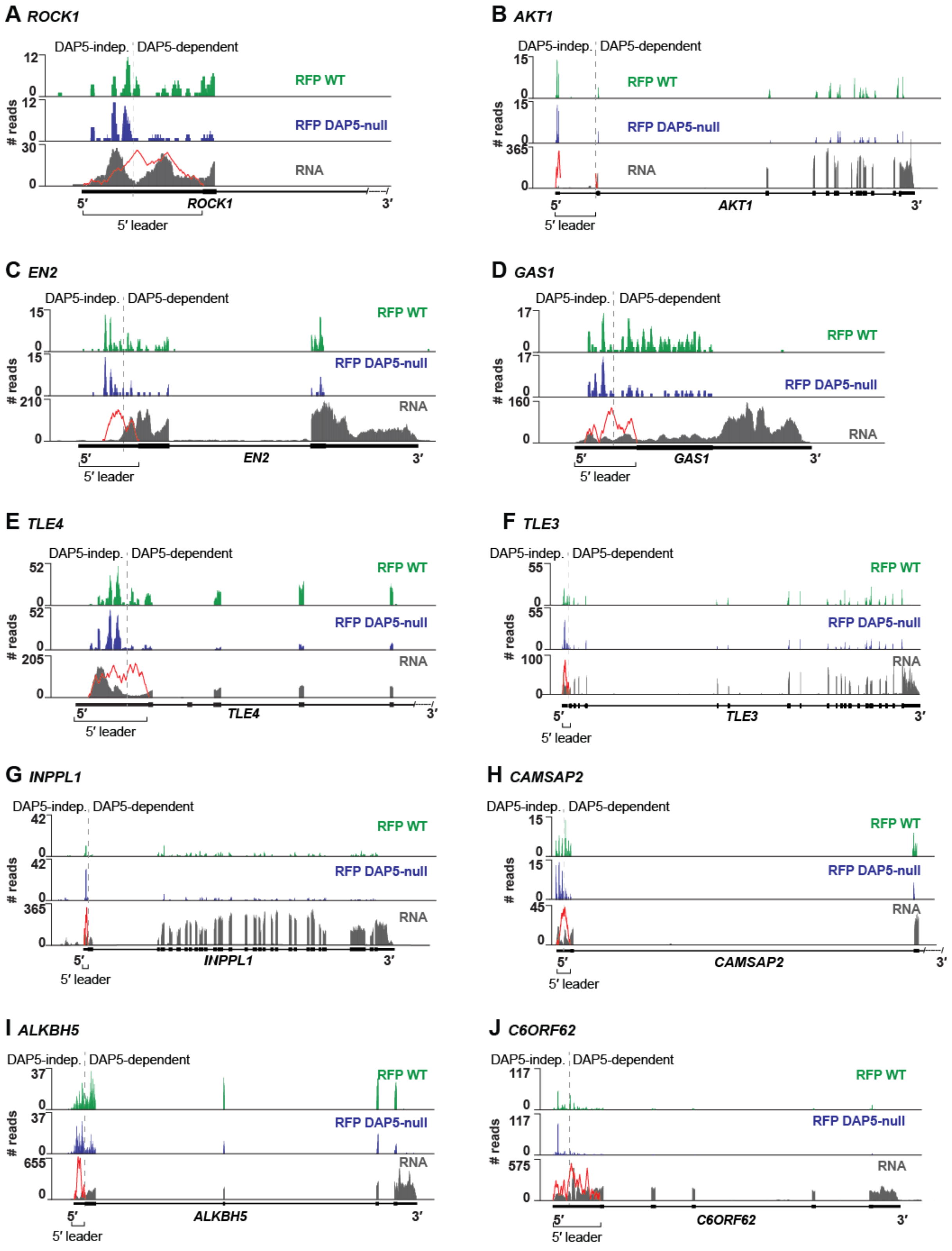


**Figure S2, related to Figure 1. Ribosome densities and mRNA read counts in DAP5 targets**

**(A-J)** Ribosome footprints and total mRNA reads distribution along the sequence of different DAP5 target mRNAs in wild type (WT) and DAP5-null cells. The predicted propensity for secondary structure across the 5′ leaders is illustrated in red. Gene annotation is depicted below the profiles. DAP5-independent (indep.) and -dependent translation is indicated with a grey dashed line.

Weber *et al.* Figure S3

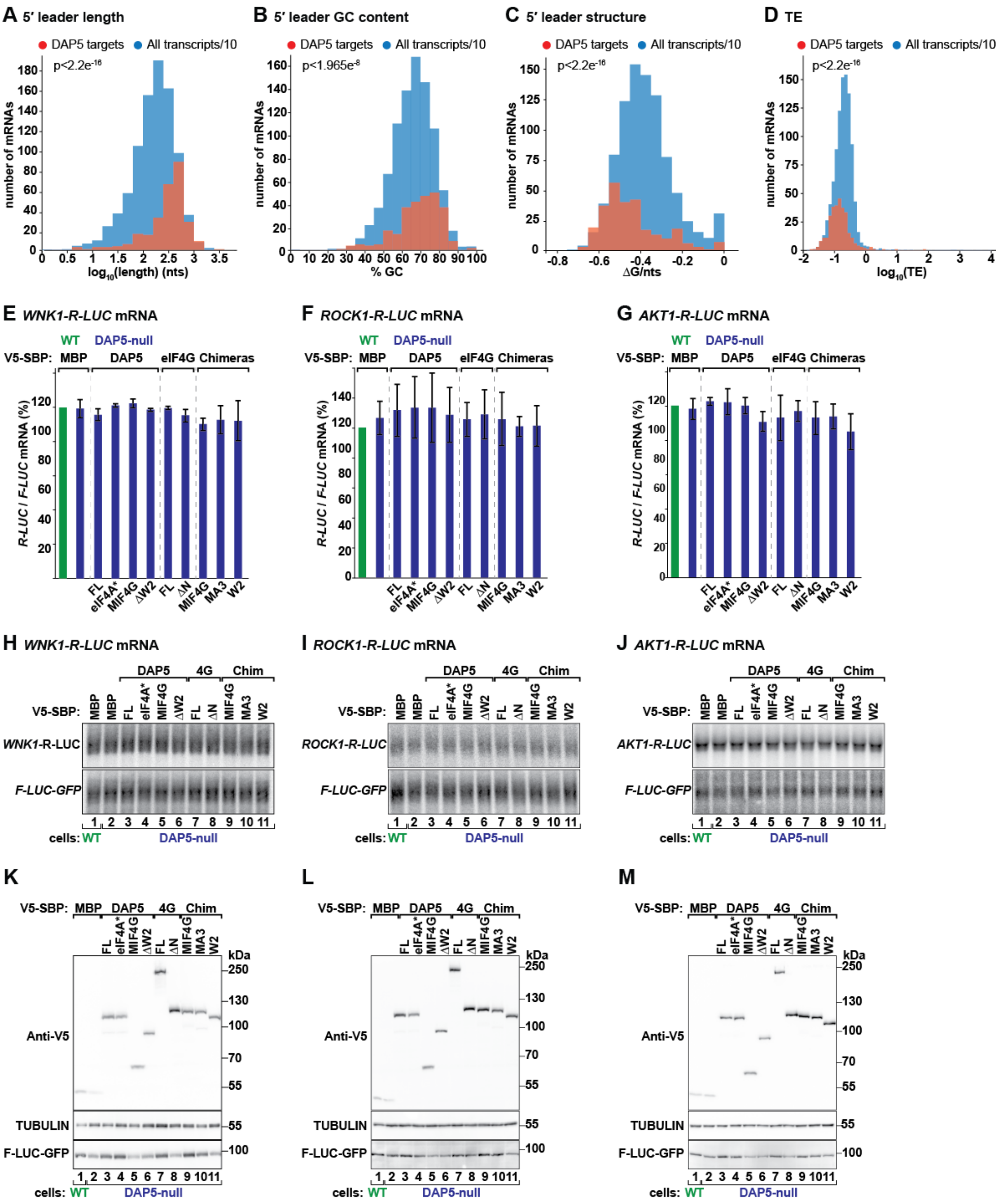


**Figure S3, related to Figures 1 and 2. DAP5-dependent translation occurs in mRNAs with structured 5′ leaders**

**(A-C)** The histograms show the distribution of length ($\log_{10}$ nts), GC content (%) and minimum free energy (ΔG/ nts) of the 5′ leaders in DAP5 targets (red) and all transcripts expressed in

HEK293T cells (blue). The number of all transcripts is reduced by a factor of 10 for visualization purposes. Statistical significance was calculated with the Wilcoxon rank sum test. Bin width is 0.2 in A, 5% in B and 0.5 in C.

**(D)** Histogram depicting the range of TE ($log_{10}$) of the DAP5 targets (red) and all other mRNAs expressed in HEK293T cells (blue). DAP5 targets show lower TE than all other transcripts ($p<2.2e^{-16}$). Statistical significance was calculated using the Wilcoxon rank sum test. Bin width=0.1.

**(E-J)** WT and DAP5-null cells were transfected with plasmids expressing *WNK1*-, *ROCK1*- or *AKT1*-R-LUC, V5-SBP-MBP, V5-SBP-DAP5 (FL or mutants), V5-SBP-eIF4G (FL or ΔN) or V5-SBP-Chimeras. *R-LUC* mRNA levels were determined by northern blotting, normalized to *F-LUC-GFP* and set to 100% in WT cells. Bars represent the mean value; error bars represent SD (n=3). Representative northern blots are shown in H-J.

**(K-M)** Immunoblots depicting the expression of the proteins used in Figures 2A-C.

**Weber *et al.* Figure S4**

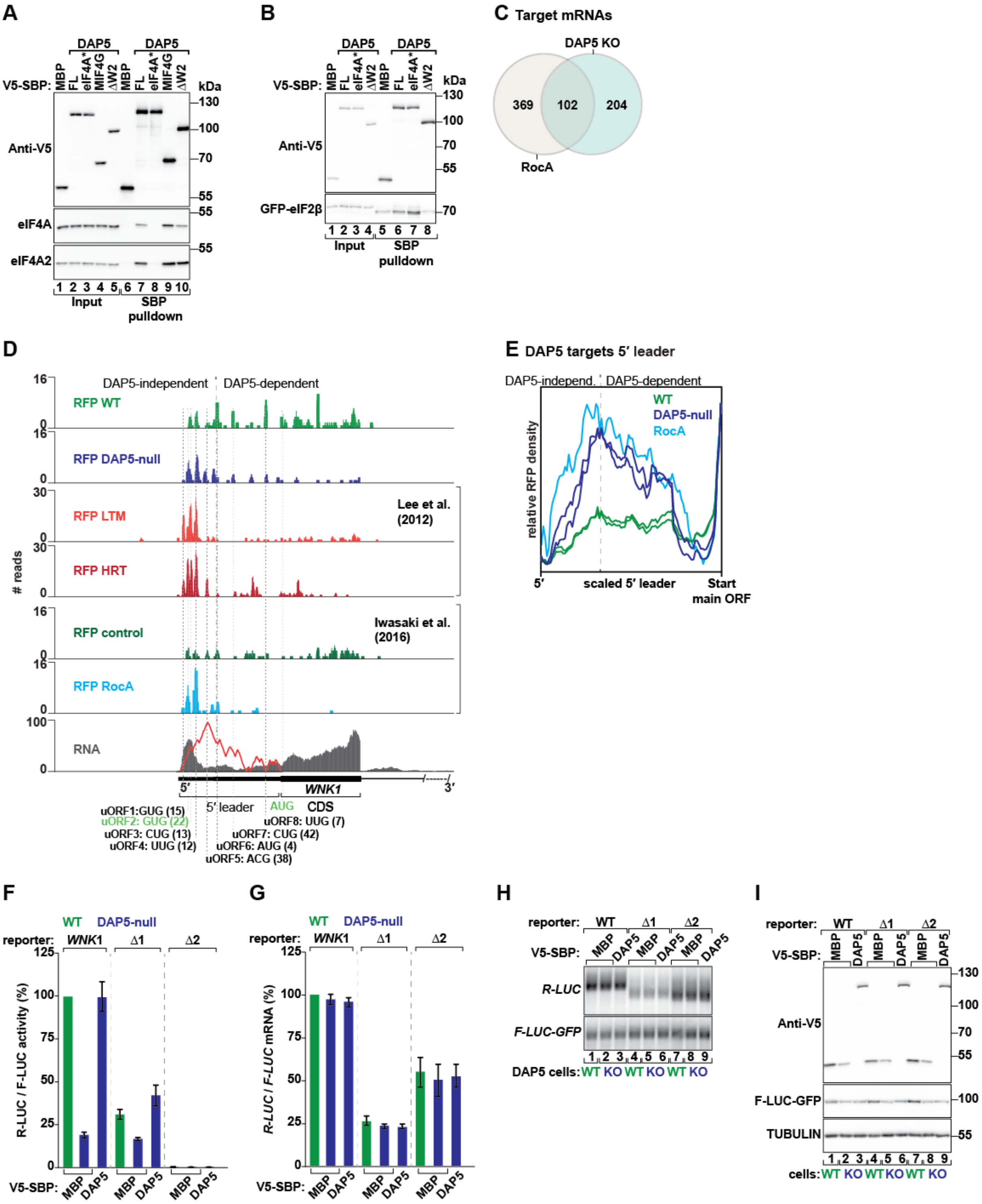


**Figure S4, related to Figures 2 and 3. DAP5-dependent translation requires the RNA helicase eIF4A**

**(A, B)** Streptavidin-binding protein (SBP) affinity pulldowns were performed two days post cellular transfection with SBP-V5-MBP or V5-SBP-DAP5 (FL or mutants) and GFP-eIF2β

(B). Input (1% for the V5 proteins, 0.3% for eIF4A, eIF4A2 and GFP-eIF2β) and pulldown fractions (1% for the V5 proteins, 2% for eIF4A, eIF4A2 and GFP-eIF2β) were analysed by western blotting with anti-V5, eIF4A, eIF4A2 and GFP antibodies.

**(C)** Venn diagram showing the number of common (n=102; p=5.1995e$^{-97}$ using a hypergeometric test) and unique genes with decreased TE in DAP5 knockout (KO) HEK293T cells and HEK293 cells treated with 0.003 μM of Rocaglamide A (RocA) (Iwasaki et al., 2016).

**(D)** Ribosome footprints and total mRNA reads distribution along *WNK1* exon 1 including the 5′ leader and the most 5′ proximal coding sequence in WT and DAP5-null cells. Also shown are the ribosome footprint profiles (RFPs) in HEK293 cells treated with harringtonine (HRT) and lactimidomycin (LTM) obtained by Lee and co-workers (Lee et al., 2012) and in HEK293 cells upon treatment with 0.003 μM of RocA (Iwasaki et al., 2016). The predicted propensity for secondary structure across *WNK1* 5′ leader is illustrated in red. uORFs position in the 5′ leader is indicated with the corresponding start codons. Start codons highlighted in green are in frame with the AUG at the main annotated coding sequence of *WNK1*. Gene annotation is depicted below the profiles. DAP5-independent and -dependent translation is indicated with a black dashed line. CDS: coding sequence.

**(E)** Metagene analysis of ribosome density for the 5′ leaders of the DAP5 targets (n=306) in WT (green), DAP5-null (dark blue), and RocA-treated cells (light blue) (Iwasaki et al., 2016). Ribosome densities were determined as the ratio of footprints within the 5′ leader relative to the footprints at the annotated downstream CDS start codon. The black dashed line indicating DAP5-independent (indep.) and DAP5-dependent translation was defined as the position along the 5′ leaders in which RFP density decreases in the absence of DAP5.

**(F-I)** WT and DAP5-null cells were transfected with plasmids expressing *WNK1*-R-LUC reporters, V5-SBP-MBP or V5-SBP-DAP5. (F) Following transfection, luciferase activities were measured (F) and mRNA levels determined by northern blotting (G, H). R-LUC activity and mRNA levels were normalized to the transfection control F-LUC-GFP and set to 100% in WT cells. Bars show the mean value and error bars indicate the SD (n=3). A representative northern blot is shown in H. (I) The immunoblot shows the expression levels of the proteins used in the assay depicted in Figure 3F. Membranes were incubated with anti-V5, GFP and TUBULIN.

Weber *et al.* Figure S5

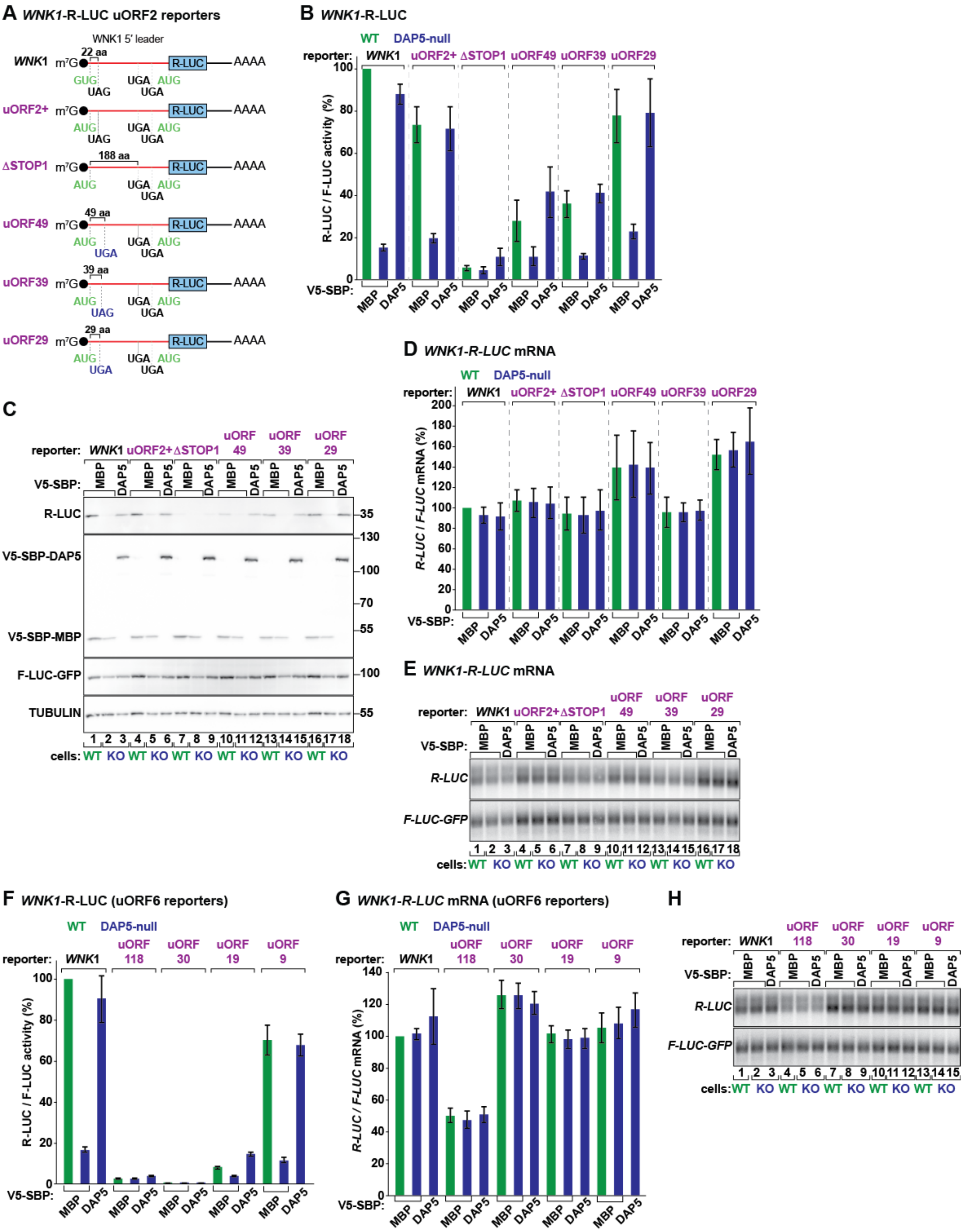

**Figure S5, related to Figures 4 and 5. Short uORFs support DAP5-dependent re-initiation**

**(A)** Schematic representations of the *WNK1*-R-LUC reporters with changes in uORF2 initiation context and length. uORF2 GUG is in frame with the R-LUC and is 22 codons long.

Three STOP codons can be found downstream and in frame with uORF2 GUG. uORF2+: GUG start codon was substituted by AUG to favour the initiation of translation. ΔSTOP1: first STOP codon in frame with uAUG was removed; uORF is then 188 codons long. uORF49, uORF39, uORF29: position of the STOP codon was moved to 49, 39 or 29 codons downstream of uAUG, respectively. aa: amino acids.

**(B-E)** WT and DAP5-null cells were transfected with different *WNK1*-R-LUC reporters, F-LUC-GFP and V5-SBP-MBP or V5-SBP-DAP5. Following transfection, luciferase activities (B) and expression (C) were measured and mRNA levels determined by northern blotting (D, E). R-LUC values were normalized to the transfection control F-LUC-GFP. The graphs show the protein (B) and mRNA levels (D) in WT and null cells, set to 100% in WT cells expressing *WNK1*-R-LUC. The immunoblot showing the expression of the different proteins is shown in panel C. TUBULIN served as a loading control. A representative northern blot is shown in E.

**(F-H)** WT and DAP5-null cells were transfected with different *WNK1*-R-LUC reporters, F-LUC-GFP and V5-SBP-MBP or V5-SBP-DAP5. Following transfection, luciferase activities were measured (F) and mRNA levels determined by northern blotting (G). R-LUC values were normalized to the transfection control F-LUC-GFP. The graphs show the luciferase activity (F) and the mRNA levels (G) in WT and null cells, set to 100% in WT cells expressing *WNK1*-R-LUC. A representative northern blot is present in H. See also Figure 5.

Weber *et al.* Figure S6

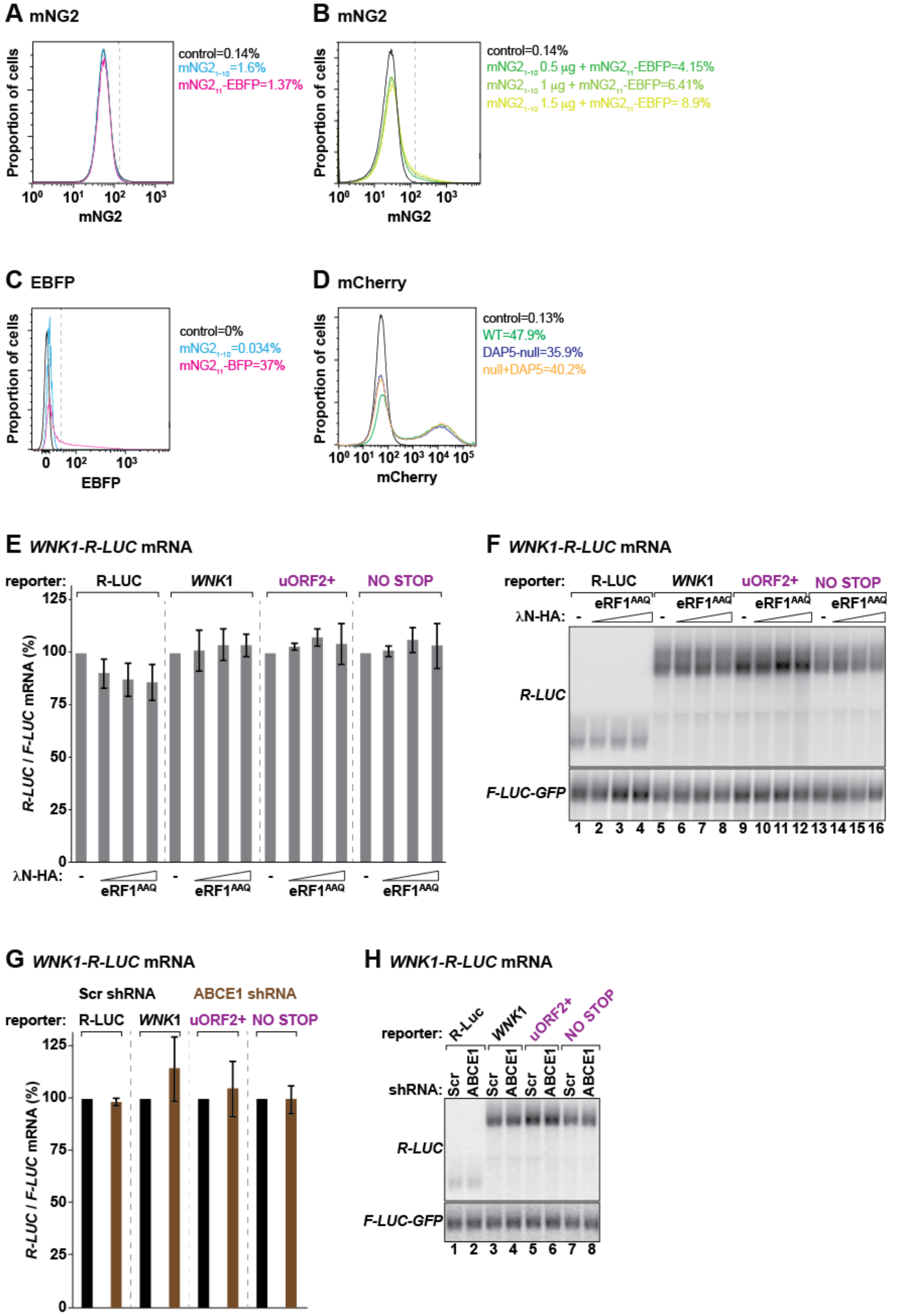


**Figure S6, related to Figures 6 and 7. Detection of uORF translation in DAP5 targets**

**(A, B)** Histograms of the mNG2 fluorescence, quantified by flow cytometry, in non-transfected (control, black trace) or HEK293T cells expressing $mNG2_{1-10}$ (light blue trace), $mNG2_{11}$-EBFP (pink trace), or increasing amounts of $mNG2_{1-10}$ (green-yellow traces) and $mNG2_{11}$-EBFP. mNG2 expression is plotted on a log scale and represents around $2x10^5$ cells.

**(C)** Histograms of the EBFP fluorescence, quantified by flow cytometry, in non-transfected (control, black trace) or HEK293T cells expressing $mNG2_{1-10}$ (light blue trace) or $mNG2_{11}$-BFP (pink trace). EBFP expression is plotted on a log scale and represents around $2x10^5$ cells.

**(D)** Histograms of the mCherry fluorescence, quantified by flow cytometry, in non-transfected cells (control, black trace), and WT (green trace) or DAP5-null HEK293T cells expressing mCherry, V5-SBP-MBP (blue trace) or V5-SBP-DAP5 (yellow trace). mCherry expression is plotted on a log scale and represents around $2x10^5$ cells.

**(E, F)** WT cells expressing increasing concentrations of $eRF1^{AAQ}$ were transfected with different *WNK1*-R-LUC reporters and F-LUC-GFP. Following transfection, luciferase mRNA levels were determined by northern blotting. R-LUC values were normalized to the transfection control F-LUC-GFP. The graph shows the luciferase mRNA levels in cells and set to 100% in the absence of $eRF1^{AAQ}$. A representative northern blot is present in panel F. See also Figure 7.

**(G, H)** Scramble (Scr) and ABCE1 shRNA-treated cells were transfected with different *WNK1*-R-LUC reporters and F-LUC-GFP. Following transfection, luciferase mRNA levels were determined by northern blotting. R-LUC values were normalized to the transfection control F-LUC-GFP and set to 100% in control knockdown cells. A representative northern blot is present in panel H. See also Figure 7.

### Weber *et al.* Figure S7

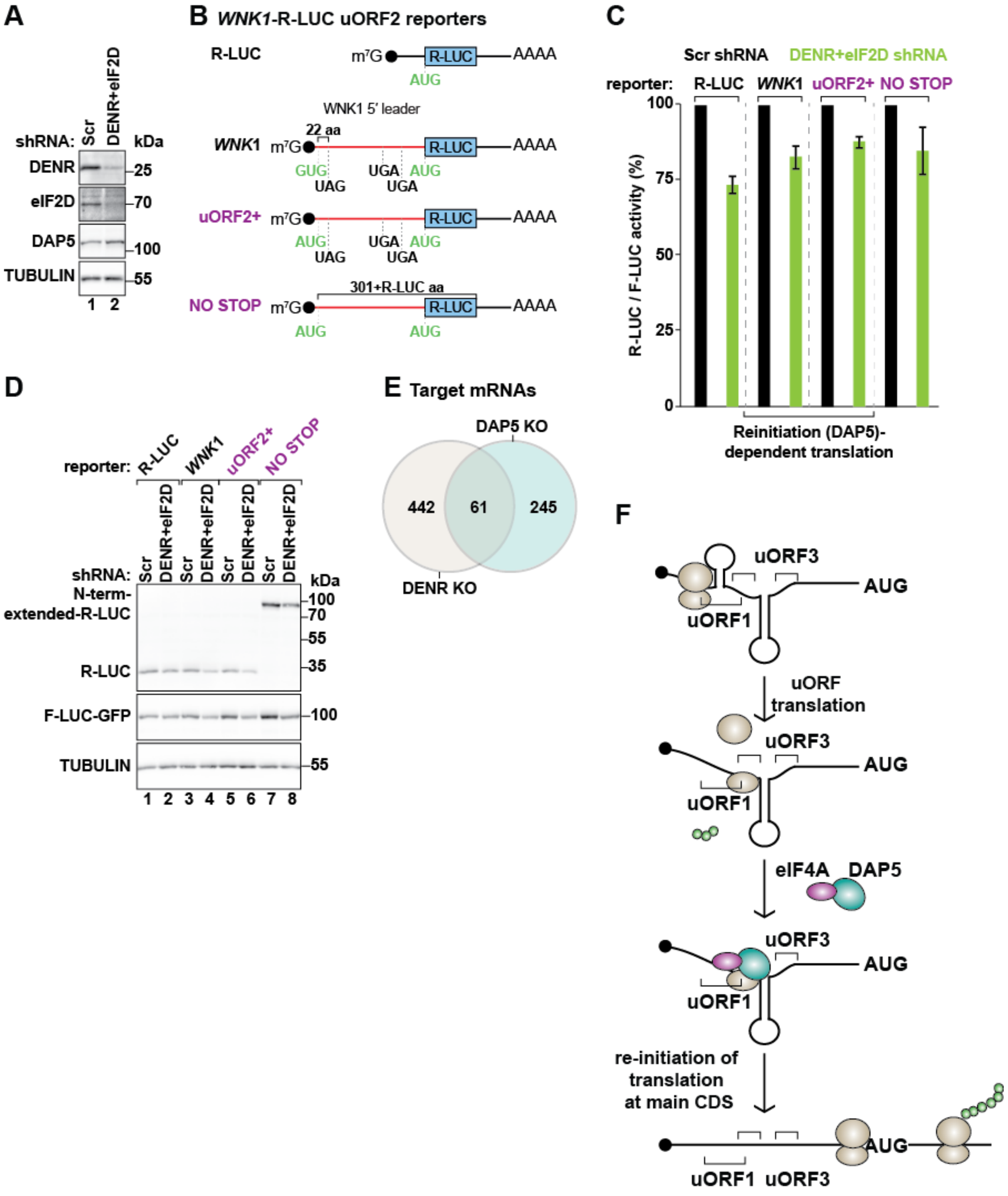


**Figure S7, related to Figure 7. Canonical translation termination and 60S recycling precedes re-initiation of translation by DAP5**

**(A)** Western blot showing shRNA-mediated depletion of DENR and eIF2D in HEK293T cells. TUBULIN served as a loading control. The levels of DAP5 were not affected in DENR/MCTS1- and eIF2D-depleted cells.

**(B)** Schematic representation of *WNK1*-R-LUC reporters with changes in uORF2 length, as described in Figure 4.

**(C, D)** HEK293T cells were treated with scramble (Scr) or shRNA targeting *DENR*, *MCTS-1* and *eIF2D* mRNAs and transfected with the *WNK1*-R-LUC reporters shown in B. The graph shows relative R-LUC activity in control (Scr) and DENR/MCTS1+eIF2D KD cells. R-LUC activity was normalized to that of F-LUC-GFP and set to 100% in Scr-treated cells for each reporter. The immunoblot illustrating the expression of short and long (N-terminally extended) R-LUC, F-LUC-GFP and TUBULIN in control and DENR/MCTS1+eIF2D-depleted cells is depicted in D. Blots were probed with anti-R-LUC, GFP and TUBULIN antibodies.

**(E)** Venn diagram showing the number of common (n=61; $p=1.2985e^{-21}$ using a hypergeometric test) and unique genes with decreased TE in DAP5 knockout (KO) HEK293T cells and DENR KO HeLa cells (Bohlen et al., 2020).

**(I)** DAP5 drives re-initiation following uORF translation. Simplified schematics of the role of DAP5 in translation. DAP5 target mRNAs contain structured regions (indicated as stem loops) and multiple uORFs (indicated as brackets) that initiate with near-cognate start codons. Secondary structures at the 5′ leaders stall PICs and trigger uORF translation. The non-optimal initiation contexts of the uORFs may result in initiation at different near-cognate start codons, as the PICs bypass some but not all uORFs (leaky scanning). Post-termination translation complexes at the STOP codon of uORFs are recognised by the DAP5 (in cyan)-eIF4A (in purple) complex which then fuels a new cycle of scanning and translation at a downstream start codon. Multiple cycles of re-initiation move the ribosome (in brown) through the burdened 5′ leaders towards the main CDS AUG. Green circles represent peptide chains. Black dot indicates the cap structure of the mRNA.

### Table S4. primers used in this study

| | | sequence (5' to 3') |
|---|---|---|
| **qPCR** | | |
| *WNK1* | fwd | CAGGGTCAGCCATCCTCAAGTA |
| | rev | GTGTCGCAACTGGGATCTGAGG |
| *ROCK1* | fwd | CCCCTCGAACGCTTTCTACAAG |
| | rev | ATCTACCTGTAGGCAAACCCGC |
| *GAPDH* | fwd | CTCTGCTCCTCCTGTTCGACAG |
| | rev | TTCCCGTTCTCAGCCTTGACGG |
| **sgRNA** | | |
| sgDAP5-a | | CACGTACCTTGGCTCGTTCA |
| sgDAP5-b | | ACACCATTGGGTTCCTCGCA |
| gDNA-locus-a1 | | GAGTTAGAACTGATCAACCAGA |
| gDNA-locus-a1 | | CCAGTATTACCTGCAGCAGGAA |
| gDNA-locus-a2 | | AGAAAGCCCTATACTAGATTCT |
| gDNA-locus-a2 | | TTCTCGCAACTCTACGGTATCC |
| gDNA-locus-b | | TTCCTGCTGCAGGTAATACTGG |
| gDNA-locus-b | | ACTTCTTCCAAAAATGGCAGAC |
| **shRNA** | | |
| Scramble | | ATTCTCCGAACGTGTCACG (Jonas et al., 2013) |
| ABCE1-1 | | GCTACAGCGAGTACGTTTACCT (Zhang et al., 2018) |
| ABCE1-2 | | CCGTGGATCTGAATTACAATT (Kara et al., 2015) |
| eIF3j-1 | | GGAATAGATGCTATGAACCCATCTT |
| eIF3j-2 | | TCGAGATGTGTGTATTTCATTGGAA |
| MCTS1 | | CAAAGGAATTGGCATTGAA (Ahmed et al., 2018) |
| DENR-1 | | GCAGATTTATCCATCGAGA (Ahmed et al., 2018) |
| DENR-2 | | GGTAATGTCAAGTGGAGTA (Ahmed et al., 2018) |
| DENR-3 | | CAAGTTAGATGCCGATTAC (Ahmed et al., 2018) |
| eIF2D-1 | | GCACAAAGAGCGTCTAATA |
| eIF2D-2 | | GAACGATCGTCATTAACTA |
| eIF2D-3 | | ACAAATGGATGAGCTGTTA |
| eIF2D-4 | | GAACTGATCAAGTCTCTGA |
| **Ribosome profiling** | | |
| 30 nt RNA marker | | AUGUACACGGAGUCGAGCUCAACCCGCAAC-P |
| 27 nt RNA marker | | AUGUACACGGAGUCGAGCUCAACCCGC-P |
| 3' adapter | | rApp/NNNNTGGAATTCTCGGGTGCCAAGG/3InvdT/ |
| 5' adapter (RNA) | | GUUCAGAGUUCUACAGUCCGACGAUCNNNN |
| Reverse transcription primer | | GCCTTGGCACCCGAGAATTCCA |
| Forward primer | | AATGATACGGCGACCACCGAGATCTACACGTTCAGAGTT CTACAGTCCGA |
| Barcoded reverse primer | | CAAGCAGAAGACGGCATACGAGATNNNNNNGTGACTGG AGTTCCTTGGCACCCGAGAATTCCA |
